# The amplitude in periodic neural state trajectories underlies the tempo of rhythmic tapping

**DOI:** 10.1101/450817

**Authors:** Jorge Gámez, Germán Mendoza, Luis Prado, Abraham Betancourt, Hugo Merchant

## Abstract

Our motor commands can be exquisitely timed according to the demands of the environment, and the ability to generate rhythms of different tempos is a hallmark of musical cognition. Yet, the neuronal basis behind rhythmic tapping remains elusive. Here we found that the activity of hundreds of primate MPC neurons show a strong periodic pattern that becomes evident when their activity is projected into a lower dimensional state space. We show that different tempos are encoded by circular trajectories that travelled at a constant speed but with different radii, and that this neuronal code is highly resilient to the number of participating neurons. Crucially, the changes in the amplitude of the oscillatory dynamics in neuronal state space are a signature of beat-based timing, regardless of whether it is guided by an external metronome or is internally controlled and is not the result of repetitive motor commands. Furthermore, the increase in amplitude and variability of the neural trajectories accounted for the scalar property of interval timing. In addition, we found that the interval-dependent increments in the radius of periodic neural trajectories are the result of larger number of neurons engaged in the production of longer intervals. Our results support the notion that beat-based timing during rhythmic behaviors is encoded in the radial curvature of periodic MPC neural population trajectories.

## Introduction

Precise timing is a fundamental requisite for a select group of complex actions such as the execution and appreciation of music and dance [1]. In these behaviors, the perception of time intervals is facilitated by the presence of a regular beat in the rhythmic sequence and individual intervals are encoded relative to this beat. This is called beat-based timing and serves as a framework for rhythmic entrainment where subjects perform movements synchronized to music [2–4]. Other set of behaviors, such as the interception of a moving target or the production of a single interval, seem to depend on a duration-based timing mechanism, in which the absolute duration of individual time intervals is encoded discretely like a stopwatch [2,5]. Functional imaging and behavioral studies have suggested the existence of a partially segregated timing neural substrate, with the cerebellum as a key structure for duration-based timing, the basal ganglia as main nuclei for beat-based timing, and MPC as potential master clock for both timing mechanisms [6–9]. Yet, the neural substrate for absolute timing, and especially for beat perception and rhythmic entrainment is still largely unknown.

Recent advances on the neurophysiology of absolute timing during single interval reproduction tasks suggest that time is represented in the structured patterns of activation of cell populations in timing areas such as MPC and the neostriatum [10–13]. Rather than being quantified in the instantaneous activity of single cells that accumulate elapsed time or encode the time remaining for an action [14–16], the duration of produced intervals depends on the speed at which the neural population response changes. This implies that the activation profiles are compressed for short and elongated for long intervals due to temporal scaling on the activity of the same population of cells [12,13].

On the other hand, MPC neurons are tuned to the duration and ordinal sequence of rhythmic movements produced either in synchrony with a metronome or guided by an endogenous tempo (synchronization-continuation task [SCT],[4,10]). Remarkably, the time varying pattern of activation of these interval-specific neural circuits follows a cascade of consecutive neural events (moving bumps) that repeats itself on each produced interval of the tapping sequence [4,10,17]. Nevertheless, single MPC cells multiplex the interval, the serial order, and task phase of the SCT, showing complex and heterogenous time-varying profiles of activation, that make it difficult to understand the neural population mechanisms behind beat-based rhythmic tapping. A successful approach to determine the latent task variables in cell populations is to project high dimensional individual neural activity into a low dimensional topological space in order to generate a robust and stable manifold [18]. Indeed, recent studies have reconstructed key hidden task parameters in the neural state population dynamics after dimensionality reduction [19–21].

Here we investigated the population dynamics of hundreds of MPC neurons in monkeys performing two isochronous tapping tasks, testing whether low dimensional state network dynamics can act as a neural clock during beat-based tapping. We found highly stereotyped neural trajectories that had two main properties during the SCT. First, the three first principal components showed a periodic path for each produced interval. Notably, these oscillatory state trajectories did not overlap across durations, a signature of temporal scaling; instead, they showed a linear increase in their radius and a constant linear speed as a function of the target interval during metronome guidance (SC), as well as during internally controlled rhythmic tapping (CC). Second, the intertrial variability of the trajectories’ radial magnitude also increased as a function of the interval, accounting for a key feature of timing behavior: the scalar property, which states that the variability of produced or estimated intervals increases linearly as a function of interval duration. These properties were highly resilient to the number of participating neurons and were replicated using simultaneously recorded cells during synchronized tapping but not during a serial reaction time control task that precluded rhythmic prediction. Finally, we found a tight correlation between the interval-associated changes in trajectory amplitude and variability during SCT, the number of neurons involved in the sequential transient activation patterns, and the duration of the neural activation periods within these moving bumps. Indeed, moving bumps simulations revealed that scaling the duration of the transient period of activity and increasing the number of neurons participating in the evolving patterns produced an increase in the radius and the variability of the corresponding neural trajectories, replicating the empirical findings. These results suggest that beat-based tapping depends of the radial amplitude of periodic state population trajectories in MPC, which depends on to the number of neurons involved and the duration of these cells’ activation periods within moving bumps.

## Results

### Rhythmic tapping behavior

We trained two monkeys (M01 and M02) in the SCT. M01 was also trained in two additional tapping tasks: the synchronization task (ST) and serial reaction time task (SRTT). During SCT the animals tapped on a push-button in synchronization with a rhythmic metronome for four times, thus producing three intervals (SC phase), followed by three internally-generated intervals (CC phase; Fig. 1A). In the ST the monkey produced five intervals guided by a metronome, similarly to the SC of SCT (Fig. 1B). Finally, during the SRTT, the animal pressed the button in response to five brief visual stimuli presented in a sequence, but separated by a random interstimulus interval, precluding the prediction of the next stimulus-response loop (Fig. 1C). Thus, during SCT and ST the animals entrained their rhythmic movements to a sensory metronome, while in the CC of SCT this was done to an internal representation of the same rhythm. On the other hand, the SRTT involved similar stimuli, tapping behavior, and sequential structure, but no predictive rhythmic timing was possible. Expectedly, the reaction times were significantly larger in the SRTT than in the ST (mean ± SD: 263 ± 37ms in the ST and 381 ± 46ms in the SRTT; ANOVA main effect of task: F(1, 718) =1443.93, p < 0.0001).

**Fig. 1.**
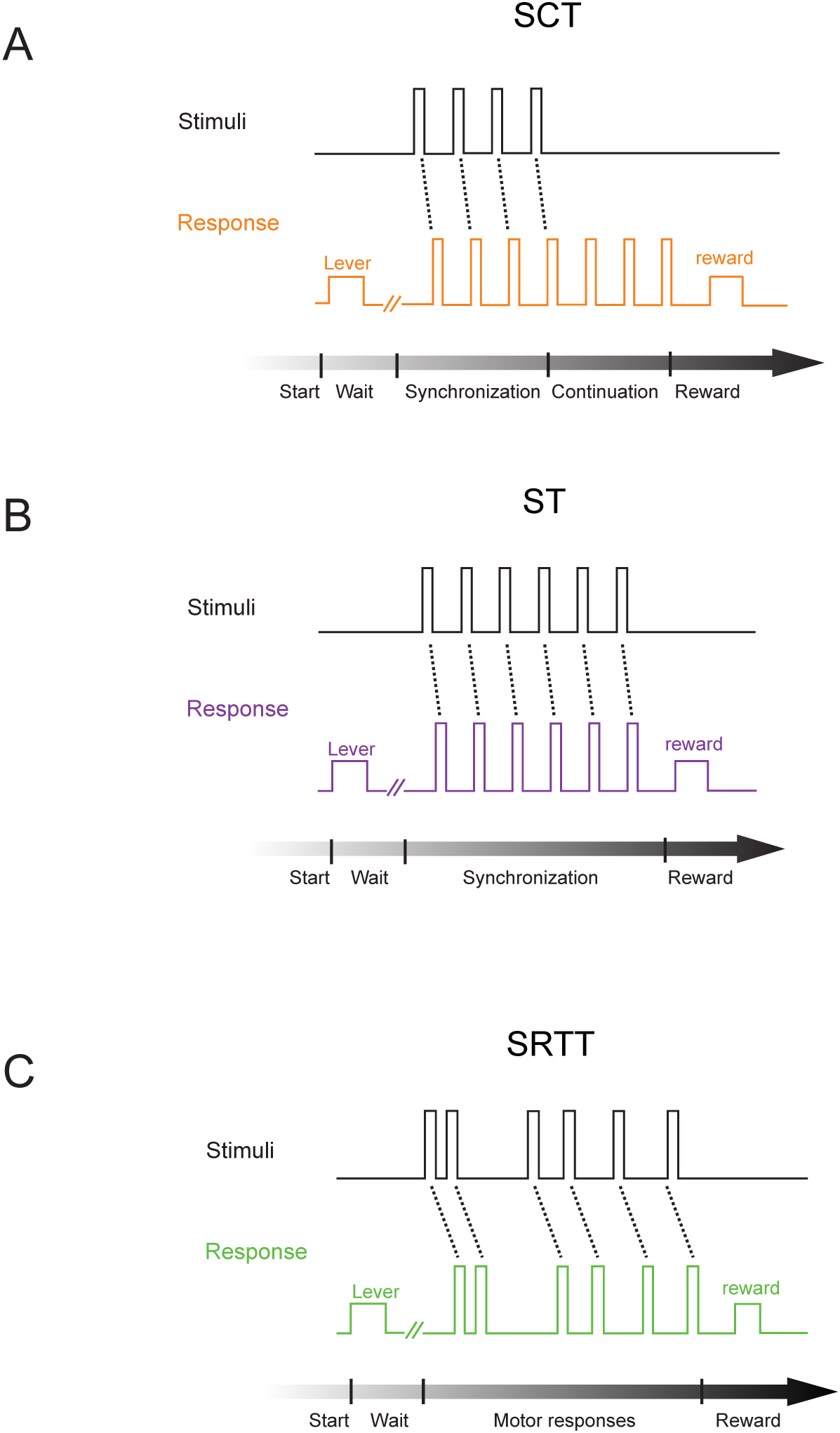
Tasks. **A**, Synchronization-continuation task (SCT). The trial started when the monkey placed his hand on a lever for a variable delay. Then, a visual metronome was presented, and the monkey tapped on a button to produce three intervals of a specific duration following the isochronous stimuli (synchronization phase), after which, the animal had to maintain the tapping rate to produce three additional intervals without the metronome (continuation phase). Correct trials were rewarded with an amount of juice that was proportional to the trial length. The instructed target intervals were 450, 550, 650, 850, and 1000 ms. **B**, Synchronization task (ST). Similar to the synchronization phase of the SCT, the animal had to produce five intervals guided by a visual metronome. The instructed intervals were 450, 550, 650, 750, 850 and 950 ms. **C**, Serial reaction-time task (SRTT). As in ST the trial started when the monkey placed its hand on a lever for a variable delay. However, in this task the monkey tapped the button after six stimuli separated by a random interstimulus interval, precluding the temporalization of the tapping behavior.

### Neural state trajectories

We characterized the dynamics of the evolving response patterns using the projection of the neural population time-varying activity onto a low dimensional state space using Principal Component Analysis (PCA) on a population of 1477 MPC cells recorded during SCT (see Methods). The results showed highly stereotyped trajectories with a strong periodicity in the first three PCs (Fig. 2A-D). Indeed, PC2 and PC3 showed together a cyclic path for each produced interval (Fig. 2C,D). Each loop in the trajectory corresponded to the periodic network state variation during the production of the rhythmic tapping sequence of the SCT. The circular trajectories in the plane exhibited the tendency to start at the same position in the phase-space after each tap, suggesting the existence of a movement-triggering point at a particular location in the population trajectory across durations (see below). Crucially, from this common phase-space location, longer intervals produced larger state trajectory loops, with a monotonic increase in the trajectory radius as a function of target interval during both the SC and CC (Fig. 2E). However, the observed interval-dependent modulations in curvilinear amplitude were not accompanied by modulations of the linear speeds of the periodic neural trajectories, as these remained constant across durations (Fig. 2F). Hence, contrary to a prototypical temporal scaling, where there is a decrease in linear speed as a function of interval and similar trajectory paths and traversed distances for different durations [13,22], the present results show that beat based-timing during the SCT is represented as an increase in curvature radii in the neural network state dynamics.

**Fig. 2.**
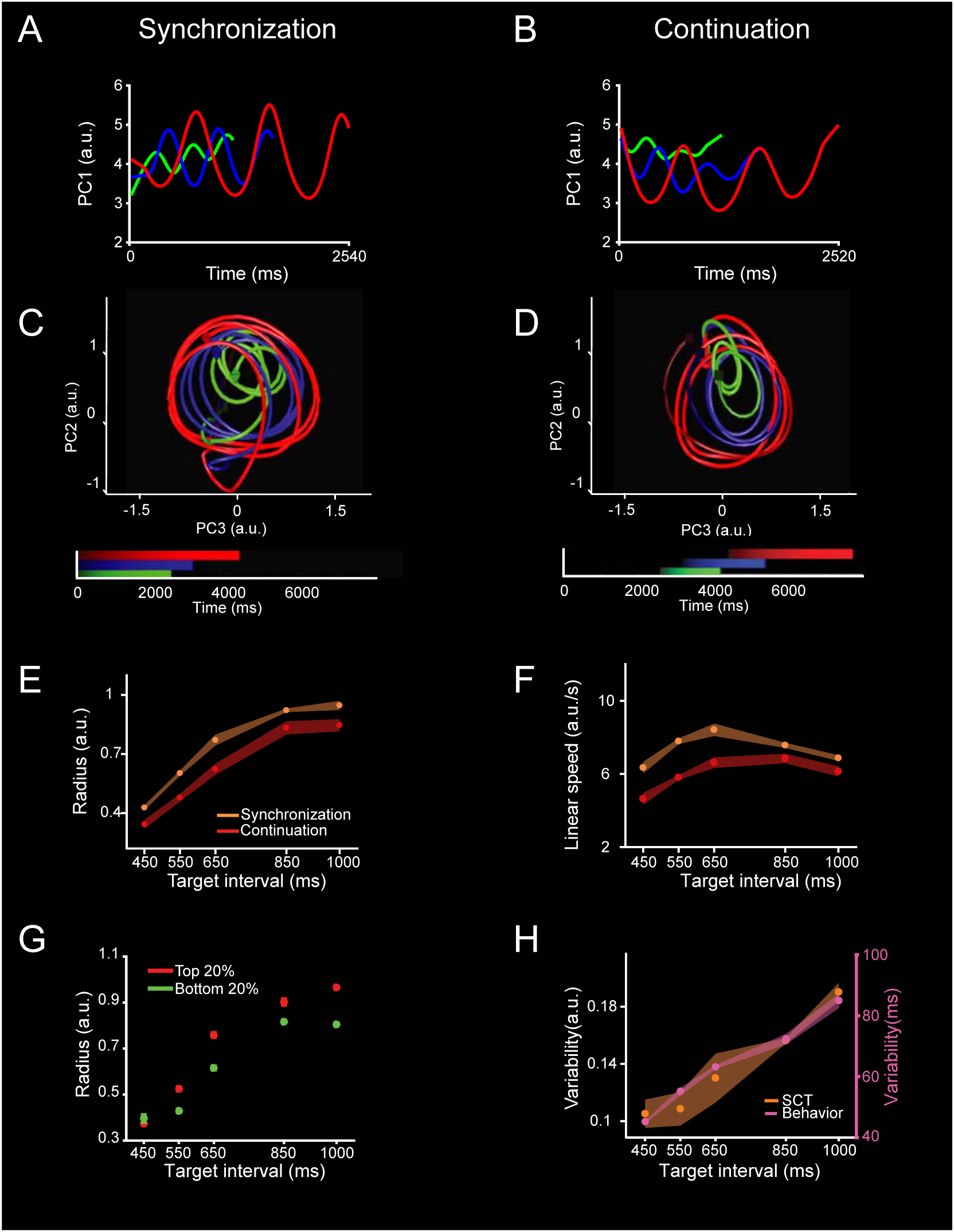
Neural population trajectories during SCT and their oscillatory dynamic properties. **A, C**, Projection of the neural activity in the MPC (1477 neurons) during the SC of the SCT onto the first (**A**) or second and third PCs (**C**). Each point in the trajectory represents the neural network state at a particular moment. The trajectory completes an oscillatory cycle on every produced interval during the synchronization and continuation phases of the SCT. Target interval in milliseconds is color-coded (450, green; 650, blue; 1000, red). Color progression within each target interval corresponds to the elapsed time. A cube marks the beginning of each trajectory, while an octahedron marks the end. **B, D**, Projection of the neural activity during CC of the SCT onto the first (**B**) or the second and third (**D**) PC. Color code is the same as (**A**). **E**, Linear increase of the Radii in the oscillatory neural trajectories during SC (red, mean±SD, slope=0.0009, constant = 0.0679, R^2^=0.9, p=0.01) and CC (orange, mean±SD, slope=0.0009, constant = − 0.0296, R2=0.9, p < 0.01) as a function of target interval. **F**, Linear speed of neural trajectories during SC (orange, mean±SD, slope=0.0001, constant = 7.322, R2=0.0007, p=0.896) and CC (red, mean±SD, slope=0.002, constant = 4.049, R2=0.354, p = 0.002) as a function of target interval (ANOVA main effect interval, F(4,39)=92.15,p<0.0001; main effect condition, F(1,39)=381.46, p<0.0001; interval x condition interaction, F(4, 39) = 15.15, p < 0.0001). The linear speed was similar (SC) or showed a slight increase (CC) with the target interval. **G,** Neural trajectory radii for the top 20% (red, slope=0.0011, constant = −0.035, R2=0.7, p< 0.0001) and bottom 20% (green,142 slope=0.00088, constant = −0.009, R2=0.75, p< 0.0001)) intertap intervals across target intervals. Note that on those intervals in which the monkeys tended to produce shorter inter-tap durations, the state trajectory radius was smaller, and vice versa (ANOVA main effect interval, F(4,40)=155.7,p<0.0001; main effect population, F(1,40)=33.3, p<0.0001; interval x population interaction, F(4, 40) = 3.98, p = 0.008). **H**, Variability (SD) of SCT rotational neural trajectories (orange, mean±SD, normalized data slope=0.0019, constant = −1.02, R2=0.94, p= 0.005) and the monkeys’ produced intervals (magenta, mean±SD, normalized data slope=0.005, constant = −0.721, R2=0.98, p= 0.0008) as a function of target interval. The Weber increase in tapping variability was not statistically different from the increase in the variability of neural trajectories across target intervals (normalized data, slope *t-test* = 0.86, p = 0.42; constant *t-test* =1.36, p = 0.22).

To test the relationship between the radius of the curvature in the neural-state trajectories and the monkeys’ behavior during SC and CC, we split the produced intervals into two groups: those in which the monkeys produced an inter-tap time that was below the 20^th^ percentile, and those with inter-tap times above the 80^th^ percentile [10]. Strikingly, on those intervals in which the monkeys tended to produce shorter inter-tap durations, the state trajectory radius was smaller, and vice versa (Fig. 2G).

Another important property of the curvilinear radii in the PCA neural trajectories was that their variability (SD of the trajectory radii) followed the same linear increase as a function of target interval observed in the monkeys’ behavior (Fig. 2H). This linear relation between temporal variability and interval duration, known as scalar property of interval timing, has been widely reported in the timing literature, and our findings suggest that it depends on the radius of the rotatory dynamical state of MPC neural populations during both SCT conditions. It is important to mention that all the described properties in the neural trajectories are resilient on the methods used to compute the PCs (see S1 Fig.).

The dynamics in the MPC population activity during the SCT was also characterized using demixed PCA (dPCA; Fig. 3; see Methods). This is a method that decomposes the dependencies of the neural population activity based on task parameters instead on the total variance explained. The first dPCA (dPCA1) showed a strong periodic structure with a minimum value around the beginning of each produced interval in the SCT sequence, similar to the findings from the PCA neural trajectories (Fig. 2C, D). In addition, the dPCA1 showed a strong change in amplitude with target duration. Since we used time-normalized neural data as input to the dPCA, all trials had the same length regardless of the target interval. In this scenario a scaling mechanism should have produced similar dPCAs across durations. Instead, we observed a time-dependent modulation in dPCA1 amplitude. In order to compare the two methods for dimensional reduction, we computed the bin-by-bin distance between the 450ms and the other four target intervals (Fig. 3F) using the PCAs (Fig. 3D) and dPCA1 (Fig. 3E). The resulting distance profiles are very similar between methods, with a periodic structure whose amplitude mean and variability increased as a function of the target interval (Fig. 3G,H). Thus, with a separate set of assumptions, the dPCA corroborates the existence of both the periodic structure of the neural state dynamics and a beat-based timing mechanism based on the amplitude modulation of the rotatory population trajectories during SCT.

**Fig.3.**
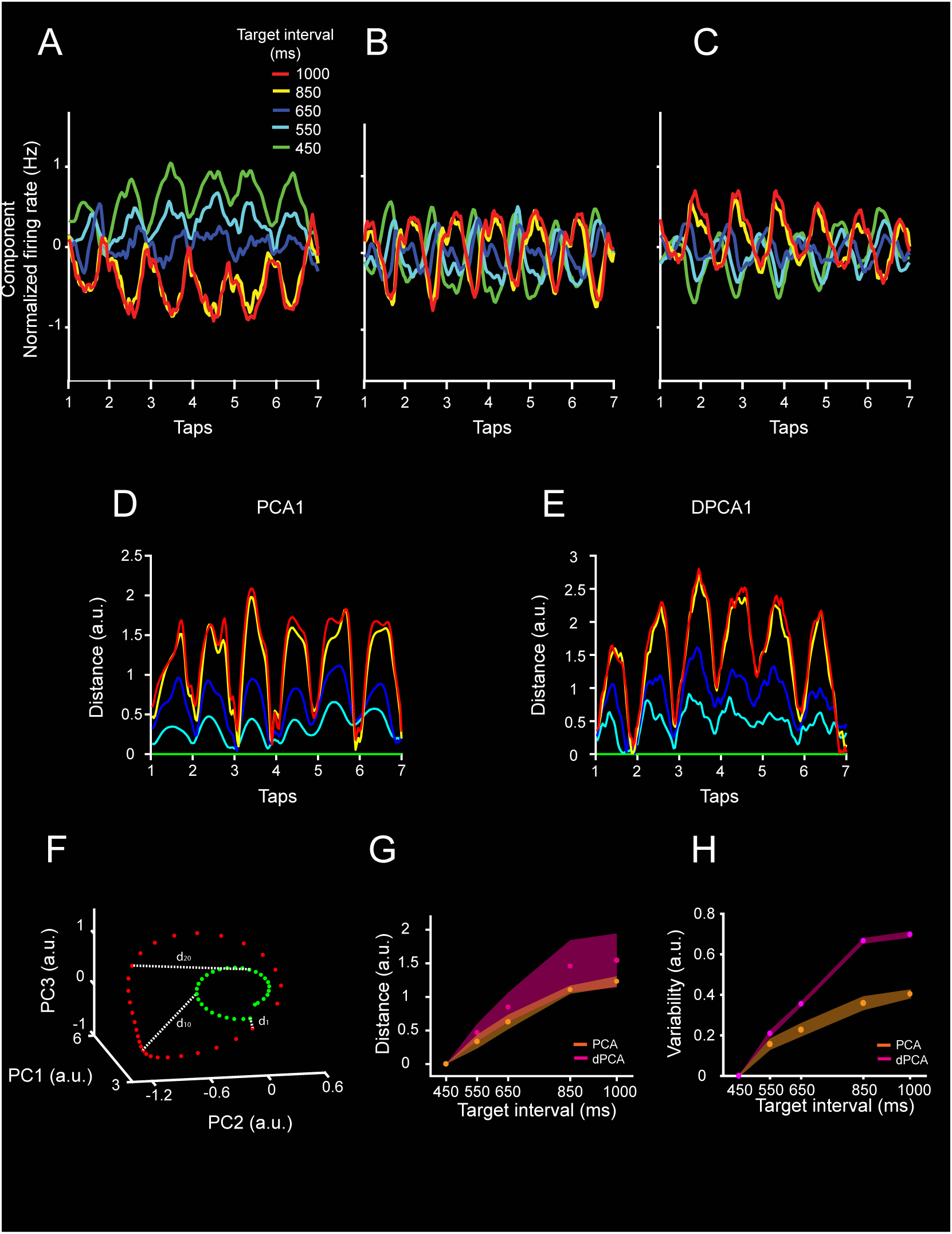
Demixed PCA applied to neural population activity during SCT. **A,B,C,** First three components of the demixed PCA of the neural activity. Together they explain 10.8% of the variance. Target interval in milliseconds is color-coded (see inset in (**A**)). Notice that the neural trajectories show oscillatory activity and their amplitude varies across target intervals. **D,E,** Euclidean distance between the first PC of the 450ms target interval and the first PC of each target interval across time for (**D**) time normalized PCA and (**E**) dPCA. Target interval is color-coded as in (**A**). Two-sample Kolmogorov-Smirnov test on the distributions of PCA and dPCA distances showed non-significant differences (p < 0.05) across target intervals. **F,** Distance calculation diagram. Binarized one inter-tap trajectories for two target intervals are shown (green, 450ms; red, 1000ms). The 450ms target interval trajectory is used as the reference for distance calculation. The Euclidean distance between each sequential bin is calculated among the reference interval and the other target intervals trajectories. Both population analysis, PCA and dPCA, produced population signals with si milar characteristics. Thus, oscillatory activity, modulation of the amplitude with the target interval, and an intersection close to the tap time are characteristics of the underlying the neural population activity irrespective of t he dimension reduction algorithm. **G,** Mean inter-tap Euclidean distance (mean±SD) between the 450ms and each target interval for the PCA (orange) and dPCA (magenta). There was a non-significant difference between the slopes of PCA and dPCA (slope *t-test* = 1.97, p = 0.0539) **H,** Variability of the distance between the 450ms and each target interval for the PCA (orange) and dPCA (magenta). The variability increased monotonically as a function of the target interval for both analysis.

The analyses described above were done on neurons recorded throughout different sessions. Thus, the neural state trajectories were also studied on simultaneously recorded cells while monkey M01 performed a synchronization task (ST, Fig. 1B) and a serial reaction time task (SRTT, Fig. 1C). As in the SCT, the PCA-projected activity during the ST showed periodic state dynamics (Fig. 4A; S3 Fig. A), whereas the SRTT neural trajectories were not as periodic (Fig. 4B; S3 Fig. B). In fact, the fitting of a normalized sinusoidal function on the first PC was statistically more robust for ST than SRTT (in terms of MSE: Fig. 4C). Again, the radius of the neural trajectories during the ST showed a significant increase in both mean radius (Fig. 4D, purple) and variability (Fig. 4E) but a constant linear speed (Fig. 4F) as a function of the target interval, reproducing the findings in SCT. In contrast, the radius and variability of the trajectories during SRTT showed small changes across target intervals, with a non-significant linear fit as a function of target interval for the three parameters (Fig. 4D, E, F green). This phenomenological comparison suggests that rhythmic tapping to a metronome depends on the amplitude of the cyclic dynamics of population activity, and that the shift from a predictive to a reactive behavior during SRTT preclude the organization of periodic population state trajectories.

**Fig. 4.**
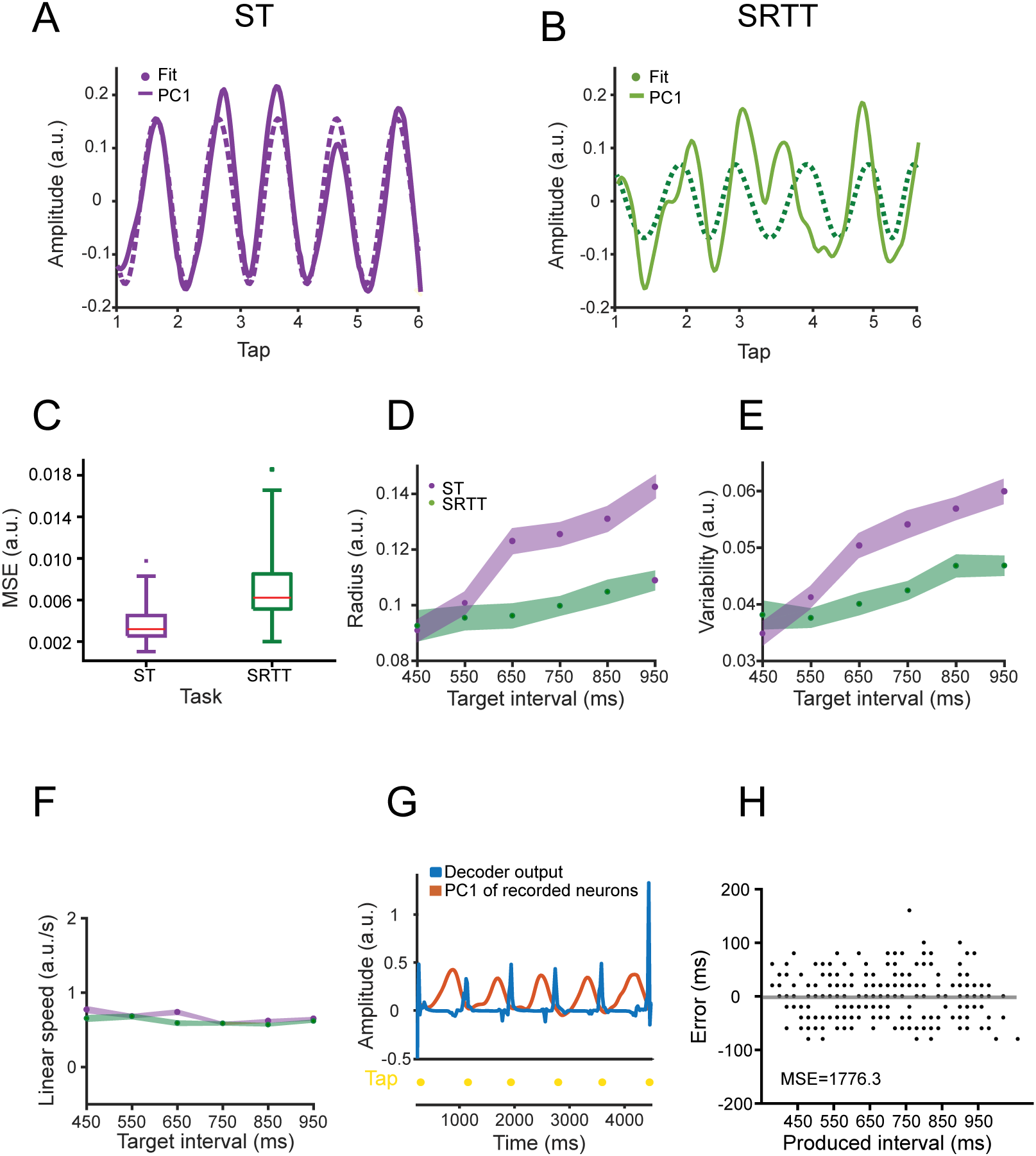
Comparison of ST and SRTT trajectories in simultaneously recorded neurons. **A**, Neural activity data projected on the PC1 (solid line, linearly detrended) and the correspondent sinusoidal fit (dotted line) during a trial of ST for the target interval of 650ms. **B**. Similar to (**A)** for SRTT. Note that the strong periodic structure of the ST neural trajectory is lost during SRTT for the same population of cells. **C**, The mean square error (MSE) of the sinusoidal fits during ST (purple) is significantly smaller than during SRTT (green; 60 trials, two-sample *t-test* = −6.78, p < 0.0001). **D**, Radii of the neural trajectories during ST (purple, slope=0.000087, constant = 0.055, R^2^=0.619, p<0.0001) and SRTT (green, **non-significant linear regression**, R^2^=0.0172 and p=0.489) as a function of target interval. **E,** Variability of the neural trajectories during ST (purple, data slope= 0.000037, constant = 0.028, R^2^=0.368, p< 0.0001) and SRTT (green, **non-significant linear regression**, R^2^=0.0005 and p= 0.903) across target intervals. **F.** Linear speed of neural trajectories during ST (purple, mean±SD, slope=0.0001, constant = 7.322, R^2^=0.0007, p=0.896) and SRTT (green, mean±SD, slope=0.002, constant = 4.049, R^2^=0.354, p = 0.002) did not change as a function of target interval. **G.** Output of the time-delay neural network (TDNN, in blue) trained to decode the duration of produced intervals based on the PC1 neural trajectories (orange) during target interval of 850ms. Tapping times are shown in yellow. **H.** TDNN error, defined as the difference between the produced and the decoded interval, as a function of produced interval. TDNN predicted accurately the performance of the monkey on a trial by trial basis (the decoded mean was not statistically different from 0, *t-test* = −0.5228, p = 0.6)

The simultaneity of the recordings during ST [23] allowed for the decoding of the produced intervals on a trial-by-trial basis. Using a time-delay neural network (TDNN, see Methods) (Fig. 4G), we found that an ideal reader of the neural trajectories could predict accurately the tapping times during ST on 86% of the produced intervals. Indeed, the decoding accuracy was better than the actual percent of correct trials in this demanding task (Fig. 4H), supporting the notion that the neural trajectories can robustly predict the rhythmic tapping behavior.

### The population state dynamics is not related to the tapping kinematics

The cyclic and smooth nature of the neural trajectories during ST and SCT sharply contrast with the kinematics of movement (Fig. 5A,C-D), that is characterized by stereotypic tapping movements separated by a dwell period that increased as a function of the target interval (Fig. 5E; [24,25]). These observations suggest that during rhythmic tapping an explicit timing mechanism in MPC keeps track of the dwell time by setting in motion a continuous and periodic change in the neural population state. According to this scheme, the tapping command is triggered once the state trajectories get to a specific position in the phase-space that correspond to the intersection point between the tangent circular paths whose radii increase with the tapping tempo. To test the hypothesis, we computed the distance between a point in state-space and the position of the taps in the neural trajectory and found a similar distance across target intervals (Fig 5B, see inset). In addition, the distance between the same point and half intertap position increased as a function of target interval (Fig 5B). Therefore, these results support the idea that the neural trajectories encode the dwell time between taps in the PC amplitude and triggers the stereotypic tapping movements once the neural dynamics reaches a point in state-space (S2 Fig).

**Fig. 5.**
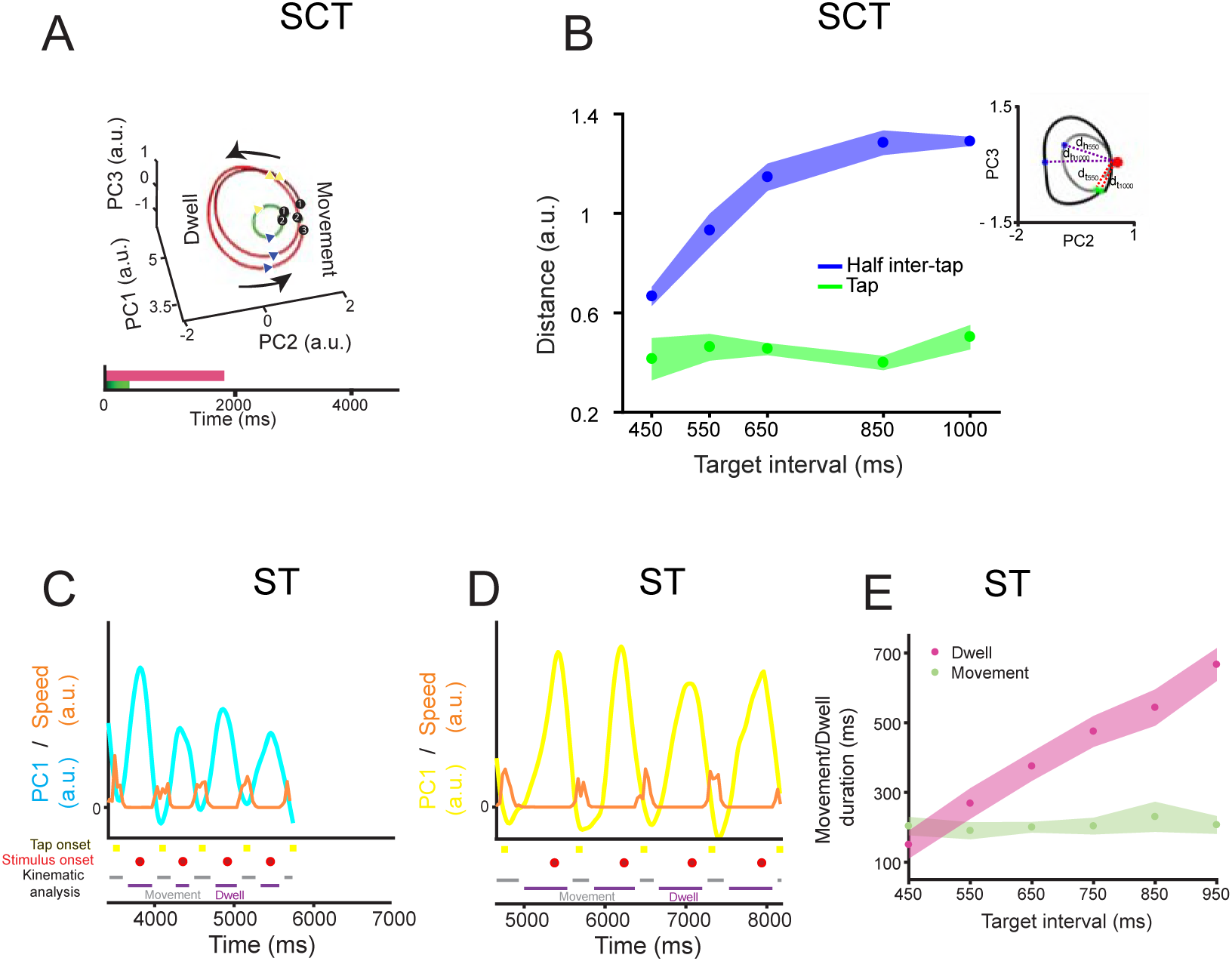
Neural trajectories do not follow the tapping kinematics. **A**, Diagram of the rotational trajectory of the SCT neural activity during three inter-tap intervals: one 450ms interval (green) and two 1000ms intervals (red). Each tap is numbered and projected in the trajectory as a white circle. A blue triangle marks the beginning, whereas a yellow triangle marks the end of the movement time. The monkeys produced phasic stereotypic movements whilst timing the dwell between taps during SCT [24]. **B,** Euclidean distance (dt, see inset) between an anchor point (red) and the position of each tap (green, mean±SD, slope= 0.00007, R^2^=0.0633, p=0.225), or half of the inter-tap interval position on the neural trajectories (blue, mean±SD, slope=-0.001, R2=0.801, p<0.<u>0001</u>) across target intervals for SC. A twoway ANOVA detected significant main effects on position (F(1,40) =1855.72, p < 0.0001), target interval (F(4,40) =77, p < 0.0001) and their interaction (F(4,40)=63.68, p<0.0001). Tukey’s HSD post hoc test showed that the distances of the anchor point to tap and half inter-tap positions were significantly different (p<0.05). In contrast, the anchor to tap distances across target intervals were not statistically different. Inset: scheme of the distance calculation, red sphere marks the anchor point, two sample inter-tap trajectories for 550ms (light gray) and 1000ms (dark gray) are shown. The green sphere marks the tap position and the blue sphere marks the half inter-tap position. Thus, the neural trajectories converge on an attractor around the tap time, to later diverge at half the inter-tap interval. Note that these results suggest the existence of tangent circular trajectories that converge in an intersection zone close to the tapping moment, although their amplitude changed as a function of interval. **C**, Speed of the tapping movement (orange trace) from the second to the sixth tap of ST, and the PC1 projected neural information (cyan) for 26 simultaneously recorded neurons during a trial with a target interval of 550 ms. Taps were represented as yellow squares and stimuli as red circles. **D**, similar to (**C**) during an 850ms target interval (PC1 projected neural information as a yellow trace). **E,** Mean ± SD of the duration of the movement (green) and the dwell between movements (magenta) across target intervals, computed from the speed profile of the tapping movements. A two-way ANOVA showed significant main effects on kinematic state (movement/dwell duration, F(1,228) =1850,61, p < 0.0001), target interval (F(5,228) = 272.72, p < 0.0001) and their interaction (F(5,228)=236.18, p<0.0001). Tukey’s HSD post hoc test showed that dwell durations across intervals were significantly different (p<0.05). Therefore, the monkey modulated the dwell duration to successfully temporalize her behavior, while the down-push-up sequence of the tapping movement was phasic and stereotypic across target intervals.

### Distributed nature of neural trajectories timing information

We determined whether we could extract information about the target interval from the neural population dynamics, and how this information was modulated by the size of the neural population used to compute the trajectories. To this end, we first segregated each segment of the single dimension trajectory according to the SCT target interval (450, 550, … 1000ms; see insets in Fig. 6A). Then, to capture the shape of the trajectory segments as a single three-dimensional coordinate, we applied a second layer PCA (PCA’) and kept the first 3 PC’s. As a result, we obtained a dot cloud in 3D where each point represents a particular produced interval trajectory segment (Fig. 6A). We trained Support Vector Machines (SVM) to classify the cloud of points for the five target intervals of the SCT. We trained the SVM ten times and used 5-fold cross validation to evaluate the performance of the classifier. On the other hand, each neuron was sorted according to the weight magnitude of the original PCAs. The neurons with the largest PC participation were removed in steps of ten percent from the original population size, and the second layer PCAs were computed on the new trajectories. Finally, the SVM was carried out on the second layer PCAs for different population sizes (see Fig. 6). There was an asymptotic decline in the classifier performance with the removal of a larger percentage of the neural population (Fig. 7A). However, even with very small populations (total cells: 15) the classifier was able to extract all SCT target interval above chance. These results are in line with the idea that the temporal structure of rhythmic behavior depends on a neural population code that is distributed within MPC.

**Fig. 6.**
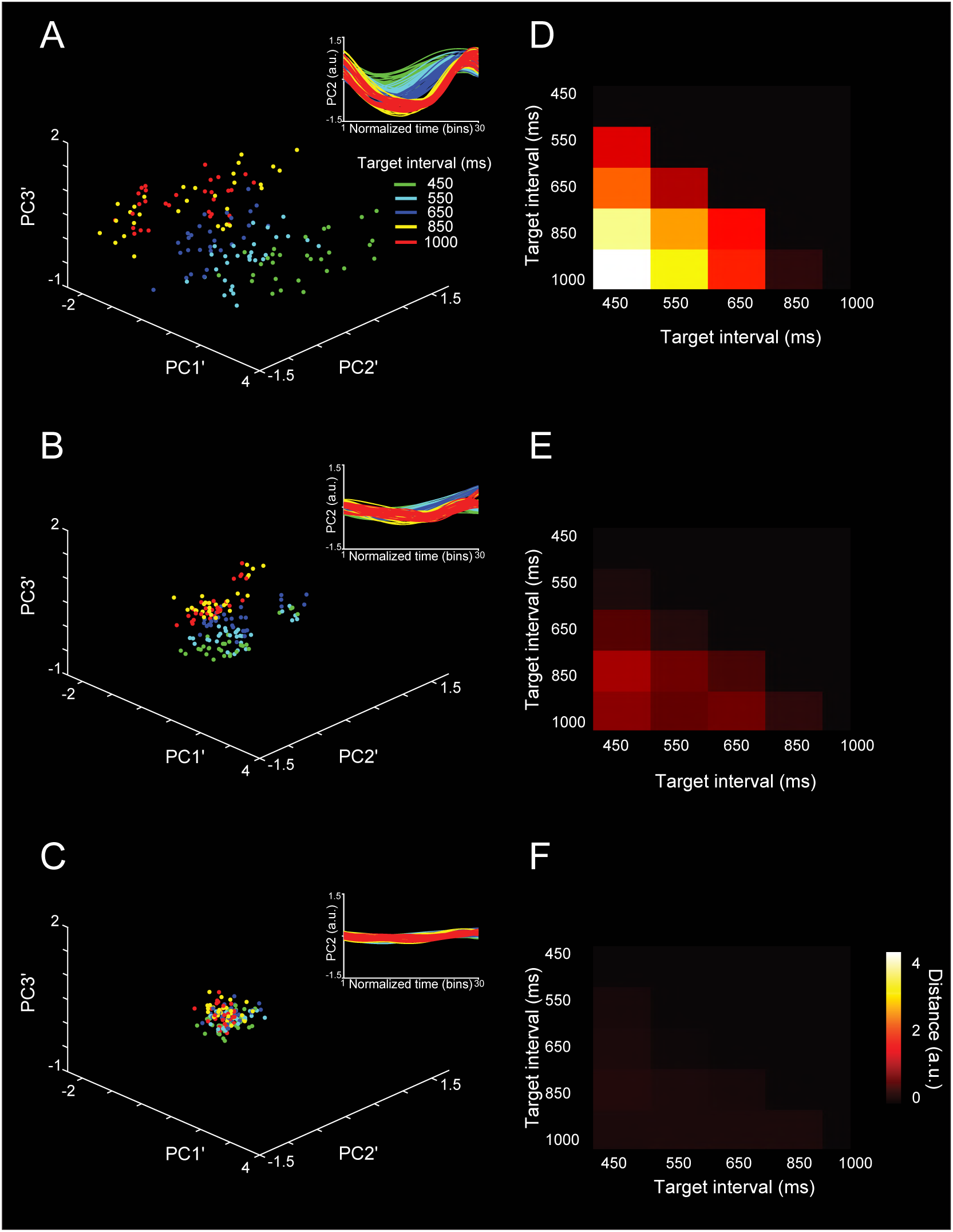
Robustness in the classifier for SCT target interval using segments of the PCA neural trajectory between taps with different neural population sizes. **A-C**, Three principal components projection of the second layer PCA applied to each of the six inter-tap neural trajectory segments and the five trial repetitions (see inset), for (**A**) 100, (**B**) 50 and (**C**) 1% of neural population. Target interval color in the inset in (**A**). **D-F**, Distances between cluster centroids of data projection across target intervals for (**D**) 100, (**E**) 50, and (**F**) 1% of neural population.

**Fig. 7.**
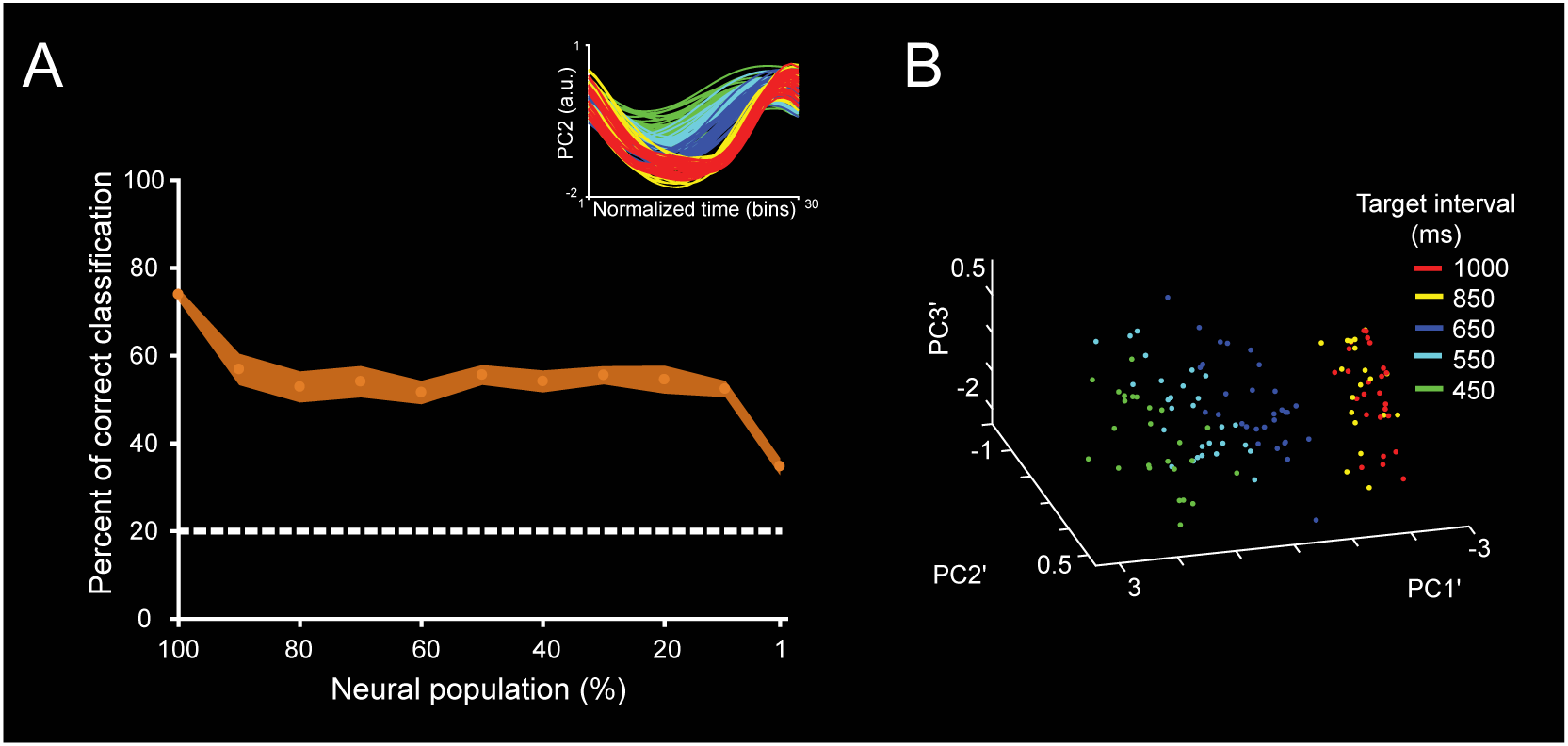
Trajectory classifier robustness across neural population sizes. **A**, Support Vector Machine classifier performance (mean ± STD of percent of correct classifications) for target interval (5 instructed intervals) during the SCT task based on the neural trajectory computed from different population sizes. The total initial population size was of 1477 neurons. Dotted lines correspond to random level. The neurons with the largest PC participation were removed in steps of ten percent of the original population size, until reaching 1% of the original population. Inset shows the original time normalized neural trajectory PC used to generate the second layer PCA’. **B**, Point cloud in 3D for the second layer PCAs’ for target interval. See color code in the inset. Note that the percent of correct classification decreased as a function of the population size; however, the classification was above chance even for the trajectories based on small cell ensembles.

### Neural population trajectories and evolving activation patterns

The results of the previous section revealed a distributed representation of tapping tempo across MPC cell populations. However, a critical question is what aspects of the time-varying activity defined the changes in amplitude in the neural trajectories as a function of the timed duration [17]. Based on our previous observations [4,10], we hypothesized that the evolving patterns of neural activity could be directly linked with the time-encoding features of the neural trajectories during the SCT. Consequently, to test this idea we first characterized the properties of neuronal moving bumps [10,12,26] during this task. With this information we carried out simulations to determine whether the key features of the moving bumps were linked to the observed changes in curvature radius and variability as a function of duration in the neural state trajectories.

As expected, a substantial proportion of MPC cells during the SCT showed a progressive pattern of activation in the neuronal population, consisting of a gradual response onset of single cells within a produced interval (Fig. 8, see Methods). This activation pattern started before a tap, migrated during the timed interval, and finished after the next tap (Fig. 8). In addition, a similar response profile was repeated in a cyclical manner for the three intervals of SC and the three intervals of CC (Fig. 8A,B) [4,10]. These findings suggest that rhythmic timing can be encoded in the sequential activation of neural populations [12]. A central question is what parameters of the neuronal response profiles are encoding the target interval and the SCT condition. Remarkably, the number of neurons involved in these evolving activation patterns (Fig. 8A,B, Fig. 9C), as well as the duration of neural activation periods (Fig. 9D) increased as a function of the target interval. SC showed a larger number of active cells whereas CC showed a longer activation period. In contrast, the neural recruitment lapse, namely, the time between pairs of consecutively activated cells (Fig. 9E), and the cells’ discharge rate (Fig. 9F) did not show statistically significant changes across target intervals and task phases. These results suggest that both the size of the circuits involved in measuring the passage of time and the duration of their activation times are core time-encoding signals in MPC, and suggest the existence of a delicate balance between these two measures to produce the progressive activation profiles of neurons when tapping to a metronome or an internally generated rhythmic signal (Fig. 9C,D).

**Fig. 8.**
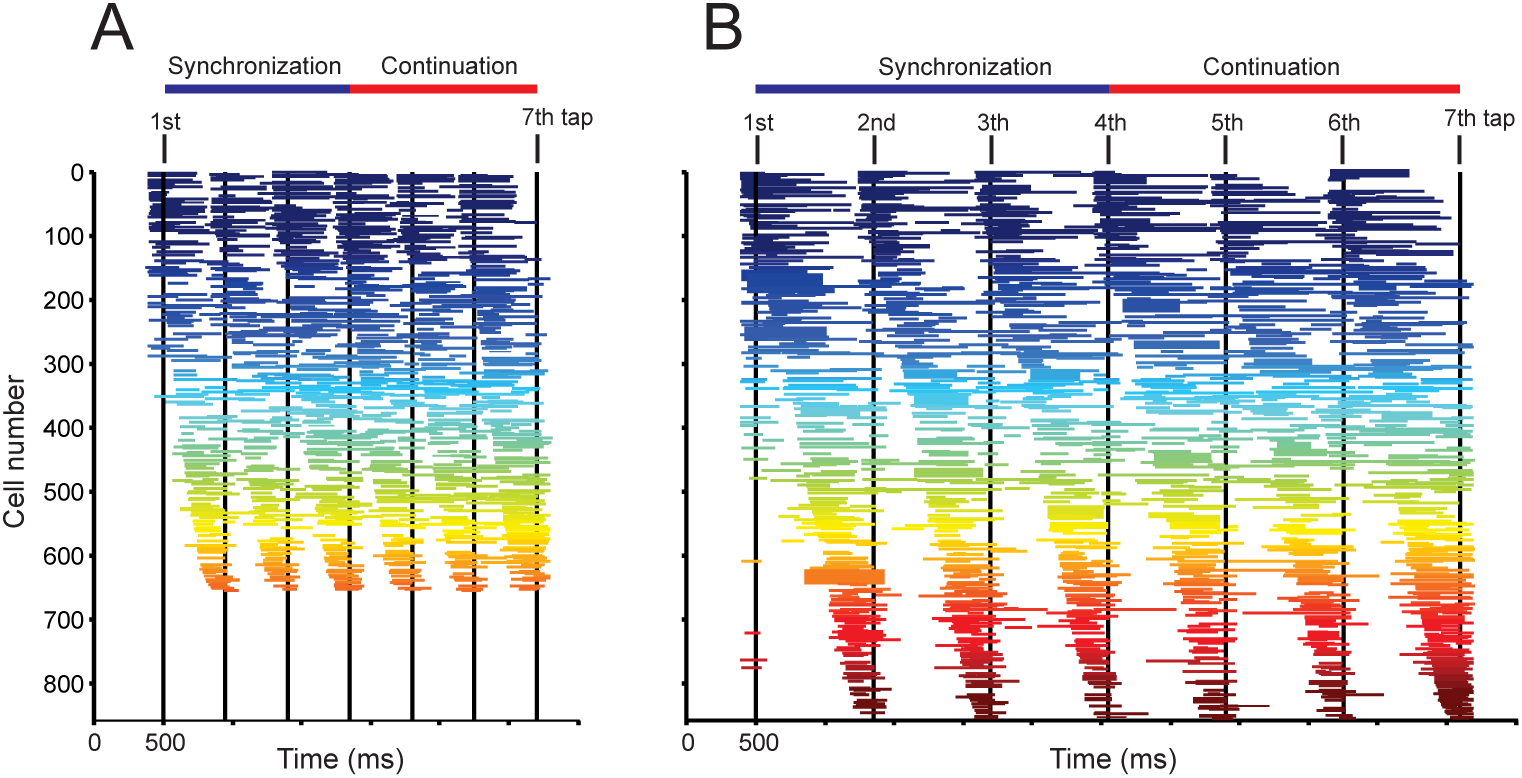
Overall patterns of activity in cell populations. **A,B,** Neural activation periods, sorted by their mean peak activation time, during the SCT task for the target intervals of 450 (A) and 850(B) ms. Each horizontal line corresponds to the onset and duration of the significant activation period of a cell according to the Poisson-train analysis (see Methods). The Poisson-train analysis was carried out on the discharge rate of cells that was warped in relation to the tapping times (seven black vertical lines; [4,49]). Note that the number of cells with significant activation periods is larger for the longer target interval.

**Fig. 9.**
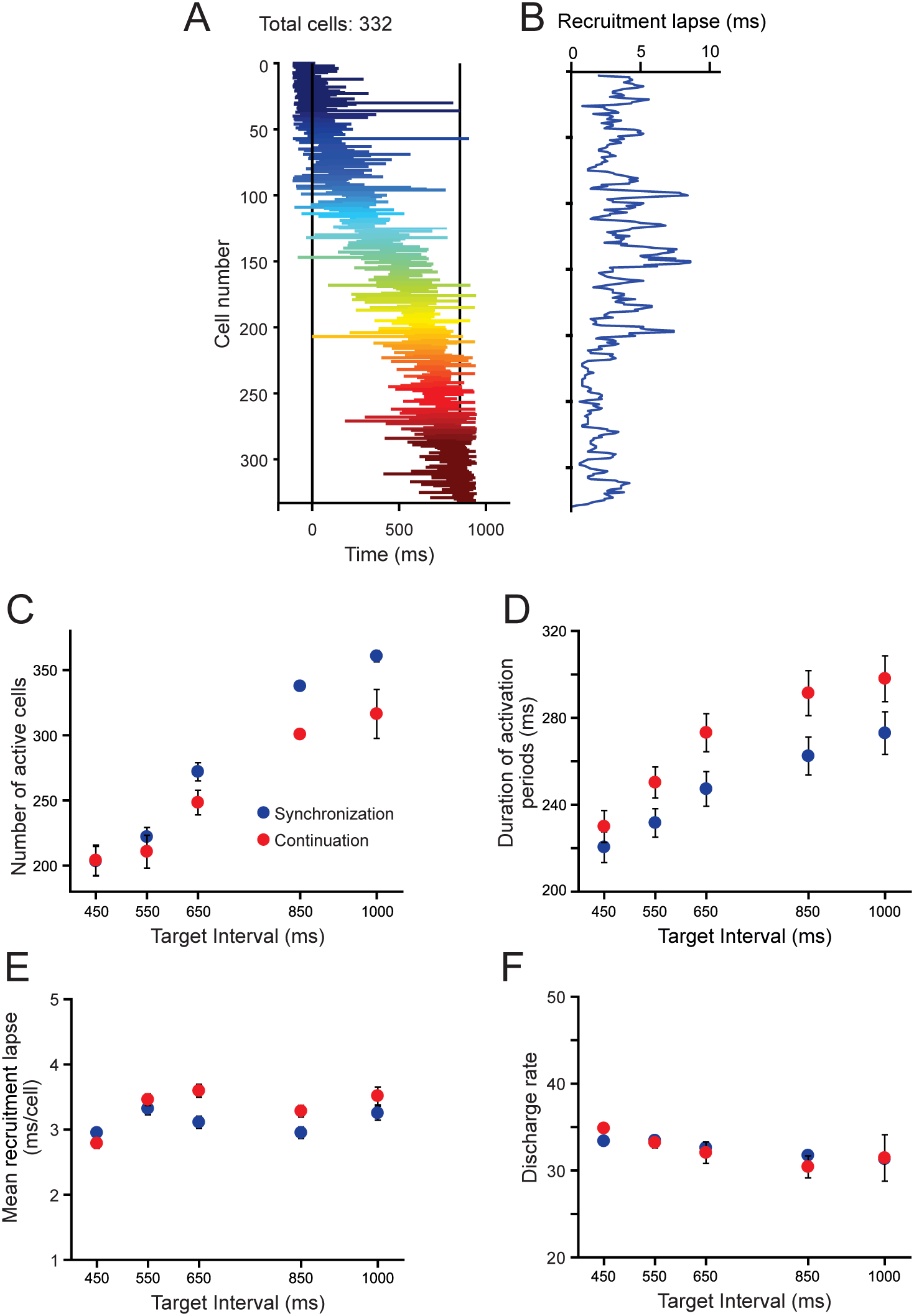
Evolving patterns of activation. **A,** Neural activation periods for the second produced interval (2^nd^ and 3^rd^ taps as black vertical lines) during SC for the target interval of 850ms. The horizontal lines of each row correspond to the onset and extent of the activation periods detected by the Poisson train analysis. Cells were sorted by their time of peak activity. **B,** Recruitment lapse as a function of cell number. The activation lapse was the difference in the time of peak activity between contiguous cells in the neural avalanche. The mean activation lapse (± SEM) was 2.98 ± 0.08 ms. **C,** Number of cells with significant activation periods across target intervals for SC (red) and CC (blue). Avalanches for longer intervals recruited more cells. (ANOVA main effect target interval, F(4,20)=21.1,p<0.0001; main effect task condition, F(1,20)=6.2, p<0.02; interval x condition interaction, F(4, 20) = 0.71, p = 0.594). **D,** Duration of the activation periods during the SC (red) and CC (blue) increased as a function of target intervals. (ANOVA main effect target interval, F(4,20)=18.9,p<0.0001; main effect task condition, F(1,20)=26.7, p<0.0001; interval x condition interaction, F(4, 20) = 1.3, p = 0.268). **E,** Mean neural recruitment lapse during SC (red) and CC (blue) did not change as a function of target interval (ANOVA main effect target interval, F(4,20)=2.7,p=0.06; main effect task condition, F(1,20)=3.4, p=0.08; interval x condition interaction, F(4, 20) = 0.79, p = 0.55). **F,** The discharge rate during activation periods in SC (red) and CC (blue) did not vary across target intervals (ANOVA main effect target interval, F(4,20)=2.2,p=0.06; main effect task condition, F(1,20)=0.86, p=0.35; interval x condition interaction, F(4, 20) = 0.92, p = 0.45).

Next, we simulated evolving patterns of population activity with different response profiles and evaluated their translation onto PCA state space. First, we generated activity patterns on individual units that were complex, heterogenous and that scaled in time, producing activation periods with the same time-varying activity but different durations (Fig. 10A, see Methods) [13]. Then, we simulated population cascade patterns for three consecutive intervals, emulating two key features on the MPC population responses: a gradual response onset of single cells that started before, migrated within, and finished after the end on an interval, with a constant overall recruitment of cells over time; and the cyclical repetition of this response profile for the three intervals (Fig. 10C,D). In addition, Fig. 11A shows that neurons were added randomly in the intermediate portion of the simulated moving bumps when increasing the total number of neurons. The projection of the simulated cascades onto PCA space produced oscillatory trajectories (Fig. 10B), whose radii and variability increased but the linear speed was similar with the target interval, as seen in the actual population responses. Importantly, these properties were only followed when the simulated neural cascades included an increase in both the number of neurons and the duration of the activation periods as a function of target interval (Fig. 10E,F). Simulations with constant values in both parameters produced PCA trajectories with similar radii or variability across interval durations, and a decrease in speed with target interval consistent with the notion of temporal scaling (Fig. 10E-G, Fig. 11B-E). Furthermore, the scaling of the response duration alone did not reproduce the observed changes in radii and variability across durations in the state trajectories (Fig. 11D-E). These findings indicate not only a close relation between the properties of the sequential neural patterns of activation and the neural state trajectories during rhythmic tapping, but also suggest that an increment in the number of neurons engaged in the evolving patterns of population activity is fundamental to reproduce the two critical duration-dependent features of the PCA neural population trajectories: the increase in the magnitude and variability of the radii as a function of target interval.

**Fig. 10.**
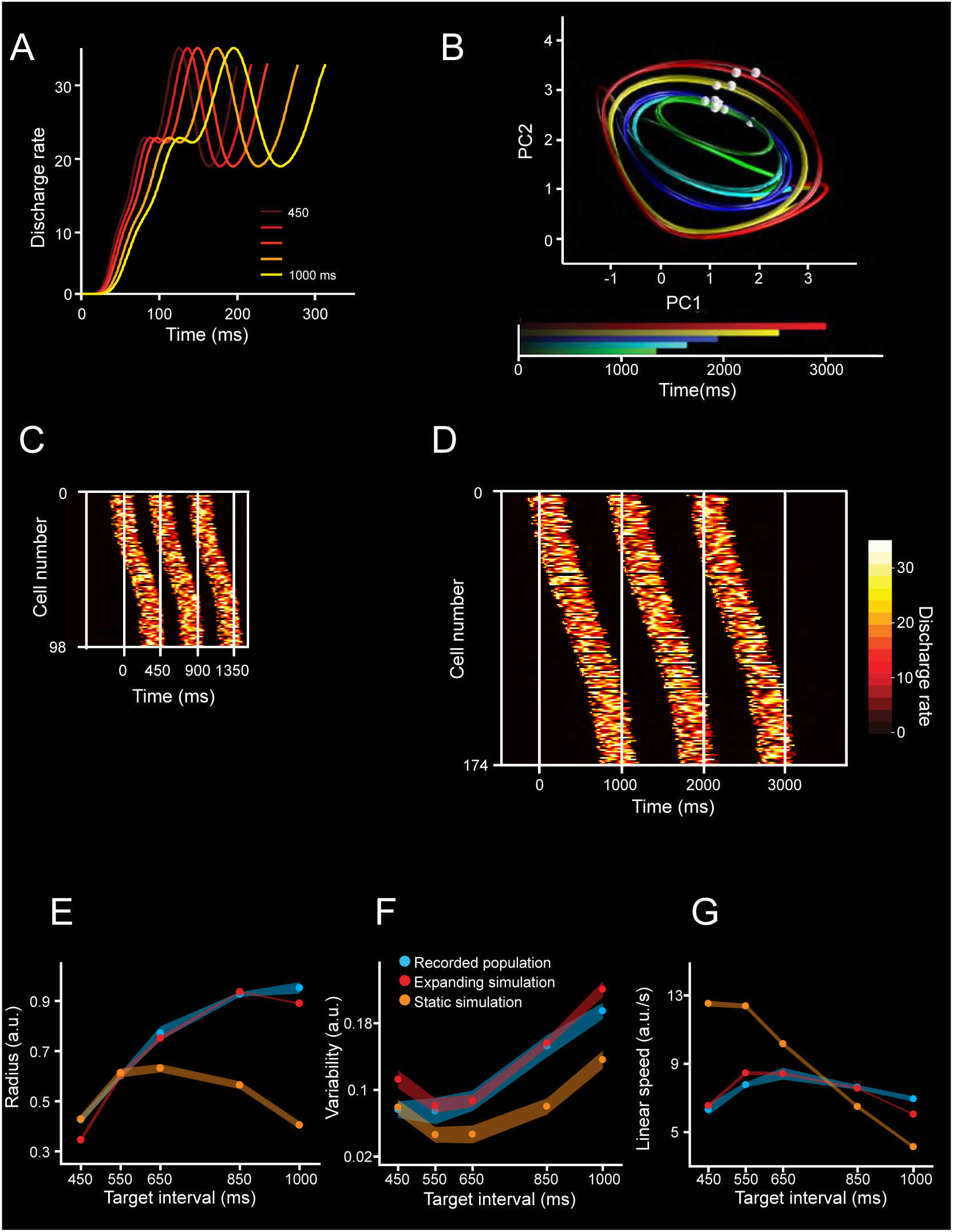
Simulations of moving bumps and neural trajectories. **A,** Activity profile of one simulated neuron during its activation period is scaled for the five simulated durations. **B**. Neural trajectories generated from the population activity of moving bumps simulations. The number of neurons and activation periods varied across intervals (see Methods). The simulated interval is color coded. Second and third simulated taps are marked as white spheres on each trajectory. **C,D**, Activation profiles of neurons for three consecutive simulated intervals with a duration of 450ms (c) and 1000ms (D). The white vertical lines correspond to the tap events defining the intervals. The activation profiles follow a Gaussian shape of cell recruitment, with slow activation rates at the tails (close to each tap). The number of neurons and the duration of the activation periods increased as a function of simulated interval. **E,F,G,** Radii (E), variability (F) and linear speed(G) of the neural trajectories generated from simulations. Data from the simulated neural activity with growing number of neurons and activation periods (red), static duration of activation periods and number of neurons (orange), and from the actual recorded population during SCT (blue) across target intervals. Note that a constant was added to both simulation data in graphs. (E) Radii for simulation with variable parameters (red, mean±SD, slope=0.0009, R^2^=0.811, p<0.0001), simulation with constant parameters (orange, mean±SD, non-significant linear regression, slope=-0.0001, R2=0.811, p=0.214), and neural activity (blue, mean±SD, slope=0.0009, R2=0.897, p<0.0001). The slopes of the radius, variability and linear speed were not statistically different between the simulations with variable parameters and the actual neuronal trajectories (radius slope *t-test* = 0.15, p = 0.878; variability slope *t-test* = 0.25, p = 0.803; linear speed slope *t-test* = 1.8, p = 0.077). However, the slopes between the simulations with constant parameters and neuronal trajectories showed statistically significant differences (radius slope *t-test* = 9.13, p < 0.0001; variability slope *t-test* = 3.73, p < 0.001; linear speed slope *t-test* = 17.71, p < 0.0001).

**Fig. 11.**
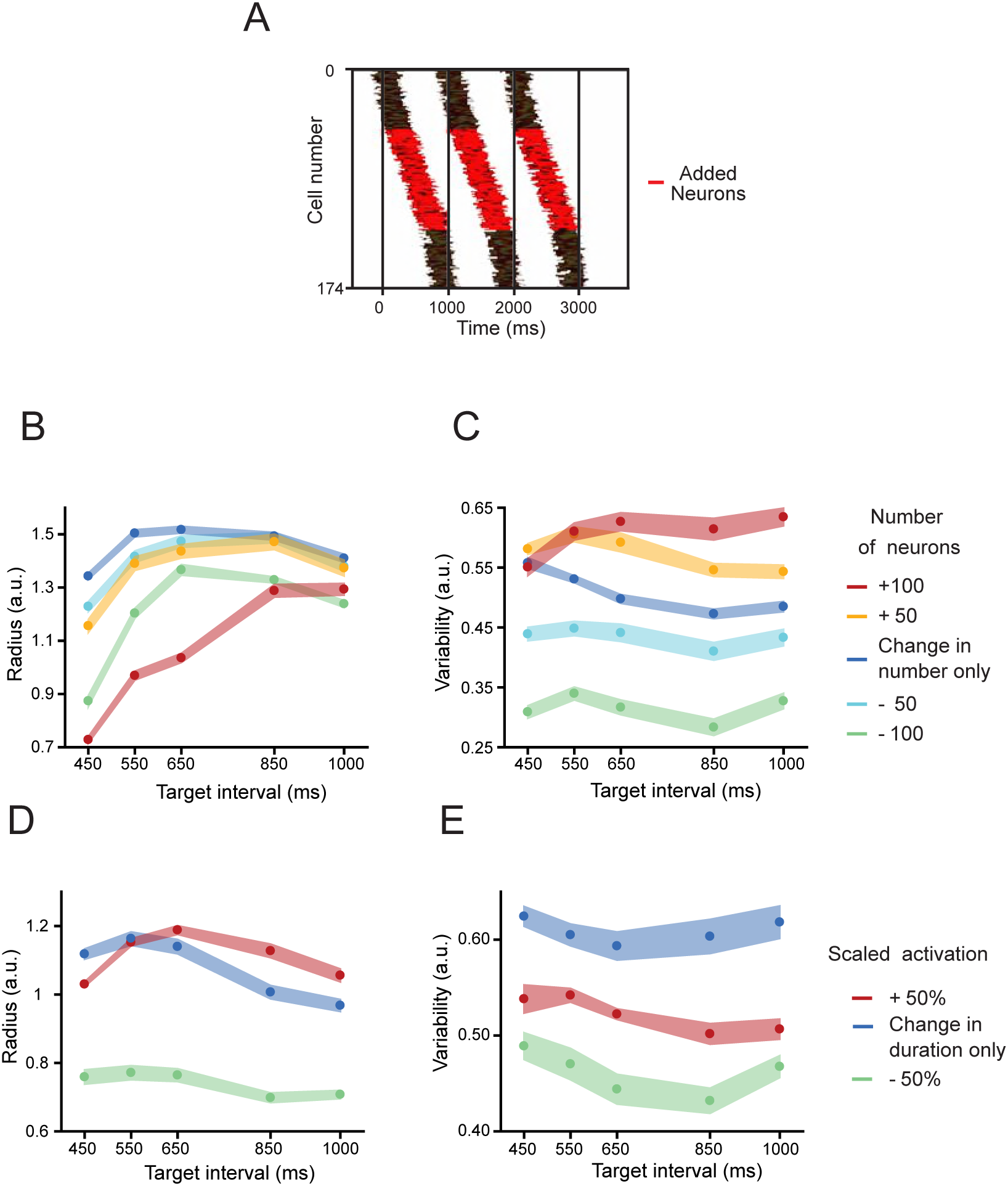
Moving bump simulation parameters. **A,** Temporal position of the activation period of neurons added to a simulation of a 1000ms target interval trial (red) in contrast to the position of the activation period of neurons also participating in a 450ms trial (black). **B,C,** Radius (B) and variability (C) of PCA trajectories generated from moving bumps simulations when the number of neurons increased a constant number of neurons below or above the original changing number of neurons as a function of target interval shown in Figure 7. A two-way ANOVA on the radius showed significant main effects for number of neurons (F(4,100) =10544. (F(4,100) = 4013.12, p < 0.0001) and their interaction (F(16,100)=25.8, p<0.0001). Tukey’s HSD post hoc test showed significant differences for the radii of all simulations with different number of neurons and for all target intervals (p<0.05). Additionally, A two-way ANOVA on the variability showed significant main effects for number of neurons (F(4,100) =2421.8, p < 0.0001), target interval (F(4,100) =3476.91, p < 0.0001) and their interaction (F(16,100)=22.53, p<0.0001). Tukey’s HSD post hoc test showed significant differences for the variability of all simulations with different number of neurons (p<0.05). **D,E,** Radius (D) and variability (E) of the trajectories generated from neural moving bumps where the duration of the activation periods was half (short, yellow) or double (long, red) than the original scaled duration (blue) as a function of target interval shown in Figure 7. A two-way ANOVA on the variability showed significant main effects for activation duration (F(2,60) =3081.54, p < 0.0001), target interval (F(4,60) =2801.16, p < 0.0001) and their interaction (F(8,60)=211.34, p<0.0001). Tukey’s HSD post hoc test showed significant differences for all simulations with different activation durations (p<0.05). In addition, a two-way ANOVA on the variability showed significant main effects for activation duration (F(2,60) =1227.53, p < 0.0001), target interval (F(4,60) =257.49, p < 0.0001) and their interaction (F(8,60)=24.87 p<0.0001). Tukey’s HSD post hoc test showed significant differences for all simulations with different activation durations (p<0.05). Thus, the number of neurons the activation duration within moving bumps produce large changes in the radius and variability of the simulated neural trajectories.

## Discussion

The present study supports four conclusions. First, the time-varying discharge rate of MPC cells shows a strong periodic organization when projected onto a two-dimensional state space, generating a circular neural trajectory during each produced interval. The amplitude of this trajectory increases with target duration and is closely related to the rhythmic tapping during the SCT and ST, but not during the reactive tapping of SRTT. Second, the scalar property, a hallmark of timing behavior, was accounted for by the variability of the curvilinear radii in the PCA neural trajectories. Third, the population dynamics for simultaneously recorded MPC cell populations during ST contained information to accurately decode the tapping times on a trial by trial basis. Last, there is a strong correlation between the interval-associated changes in radial magnitude and variability of the periodic neural trajectories during SCT and the number of neurons involved in the sequential activation patterns as well as the duration of their transient periods of activation within these moving bumps.

The network state trajectories showed the following properties: they were simple, periodic, exhibited an amplitude modulation according to the timed duration, and were different from the stereotypic kinematics of the phasic tapping movements and the timing control of the dwell between movements in this task [24,25]. Notably, the increases in trajectory amplitude as a function of target interval were observed during the two rhythmic tapping tasks, reproduced with dPCA, and closely related with the monkeys’ produced intervals during SCT and ST. Furthermore, the switch from predictive rhythmic tapping to a reaction time task (SRTT) produced a profound disorganization in the periodicity of neural trajectories accompanied by no changes in radial amplitude. In contrast with the temporal scaling model [13] we found that the neural trajectories do not scale in time, because they present a time-related amplitude modulation with similar linear speed profiles across durations. In line with our observations, neural-network simulations of complex sensorimotor patterns showed that temporal scaling of input stimuli produced curvilinear trajectories that increased in radii for longer intervals [27]. Hence, temporal scaling may be associated with interval-based timing whereas amplitude modulations in neural population trajectories can be associated with beat-based timing [28] or complex temporal processing [27].

We found a strong correlation between the duration of the produced intervals and the curvilinear amplitude of the MPC neural trajectories during the SCT and ST and, due to the simultaneity of the recordings in the latter task, we decoded accurately the produced durations on a trial by trial basis. In addition, the cyclic and smooth nature of the neural trajectories during ST and SCT sharply contrasts with the tapping kinematics, which are characterized by stereotypic tapping movements separated by a dwell period that increases with the timed interval [24,25]. Previous studies have demonstrated that cell populations in premotor and motor cortical areas show rotatory non-muscle like trajectories that reflect the internal dynamics needed for controlling reaching and cycling [29,30]. Under this scenario, we found evidence supporting the notion that the periodic MPC trajectories during rhythmic tapping encode the dwell between taps in their curvilinear radii and that the tapping command is triggered whenever the trajectory reaches a specific phase-space, which corresponds to the intersection point between the tangent circular paths. This dynamical geometry contrasts with the neural trajectories of medial frontal areas during a single interval reproduction task [22]. In this interval-based paradigm the state trajectories not only evolve at different speeds but also generate parallel paths for different timed intervals depending on the initial conditions of the neural population dynamics [22]. Thus, the present data are consistent with the notion that timing is encoded in a neural population clock [17,31–34] and puts forward the hypothesis that temporal processing during beat entrainment depends on the amplitude of tangent circular trajectories in MPC populations.

The scalar property states that temporal variability increases linearly as a function of timed duration [35]. This hallmark feature of temporal processing has been documented across many timing tasks and species [9,35–38]. Several computational models based on neural population time representations have been implemented to describe this property including drift diffusion [39,40] and recurrent networks [26,41]. Here, we found that the variability in the radii of neural trajectories increased as a function of target interval during SCT and ST but remained similar during the SRTT, a task that precludes time prediction while preserving the sensory and tapping components. Therefore, these results suggest that the amplitude of the MPC state-network trajectories is a feasible neural correlate of the scalar property during rhythmic tapping.

The dynamics of coordinated neural population activity define the evolution of the network state trajectories, which in turn have revealed functional principles in a variety of behaviors that are not evident at the single cell level [13,19,21,42]. Notably, the tapping tempo is strongly mapped in the neural trajectories and is encoded in a distributed fashion, not dependent on a particular response profile of individual neurons. Within this neural population framework, we found large groups of neurons that showed sequential transient activation patterns that traversed each produced interval during the SCT. Previous studies have reported moving bumps as a timing mechanism in parietal cortex [43]; MPC [4,10], the basal ganglia [12,44,45], and hippocampus [46,47]. For example, the bump activity in the rat striatum during a peak interval task moved progressively slower as the timed interval progressed, providing a functional basis for the decrease in the animals’ timing accuracy as the length of the timed interval increased [12]. In contrast, during the SCT we found that the rate of engagement of the neurons within moving bumps was constant and was accompanied by an increase in the number of neurons participating in the evolving patterns of population activity. Thus, an optimal reader could estimate the tempo of rhythmic tapping based on two signals: the location of the activity within a bump, where longer intervals engaged moving bumps composed of larger number of neurons, and the resetting between consecutive evolving activation patterns [40]. Strikingly, our simulations revealed a tight relation between the scaling of the duration of the transient period of activity, the increase in the number of neurons within moving bumps, and the increase in radius and variability of the corresponding neural trajectories. The simulations also suggest that neurons have the same relative position within a moving bump independently of the timed interval, as seen previously in the rat striatum [12]. Consequently, the increase in neural population size for longer intervals implies that incorporation of new cells at intermediate locations within the moving bump [10]. These results not only replicate our empirical observations, but also support the notion that the properties of moving bumps, especially the number of participating neurons, can shape the curvilinear amplitude and the corresponding variability in neural state trajectories during SCT.

Overall, these findings support the notion that the beat-based mechanism for rhythmic tapping is based on the changes in curvature radii of the neural population state dynamics in MPC, with slower tempos encoded in larger traversed distances in the tangent periodic neural trajectories and suggest that the variability in these neural trajectories is a feasible neural substrate of the scalar property during rhythmic tapping.

## Materials and Methods

### Subjects

All the animal care, housing, and experimental procedures were approved by Ethics in Research Committee of the Universidad Nacional Autónoma de México and conformed to the principles outlined in the Guide for Care and Use of Laboratory Animals (NIH, publication number 85-23, revised 1985). The two monkeys (M01 and M02, *Macaca mulatta*, both males, 5-7 kg BW) were monitored daily by the researchers and the animal care staff to check their conditions of health and welfare.

### Tasks

#### Synchronization-Continuation Task (SCT)

The SCT has been described before [48]. Briefly, the monkeys were trained to push a button each time stimuli with a constant interstimulus interval were presented. This resulted in a stimulus-movement cycle (Fig. 1A). After four consecutive synchronized movements, the stimuli were eliminated, and the monkeys had to continue tapping with the same interval for three additional intervals. Monkeys received a reward (drops of juice) if each of the intervals produced had an error < 30% of the target interval. The daily performance of the monkeys was >70% of correct trials. The amount of juice was proportional to the trial length. Trials were separated by a variable intertrial interval (1.2-4 s). The target intervals, defined by visual stimuli (red square with a side length of 5cm, presented for 33ms), were 450, 550, 650, 850, and 1,000 ms. The target intervals were chosen pseudorandomly within a repetition. Five repetitions were collected for each target interval.

#### Synchronization Task (ST)

This task was similar to the synchronization phase of the SCT [25]. The subject had to push a button with a stimulus. Six stimuli with a constant interstimulus were presented (red square with a side length of 5cm, shown for 33ms). Thus, the metronome was always present during the task. The target intervals were 450, 550, 650, 750, 850, and 950ms. Five repetitions were collected for each target interval.

#### Serial Reaction Time-Task (SRTT)

This task is also described elsewhere [48]. Monkeys were required to push a button each time a stimulus was presented, but in this case the interstimulus interval within a trial was random (picking randomly from the same 450,550,650,750,850, or 950ms), precluding the explicit temporalization of tapping (Fig. 1B). Monkeys received a reward if the response time to each of the five stimuli was within a window of 200 to 500 ms. The intertrial interval was as ST. Visual (white square with a side length of 5cm, presented for 33ms) stimuli were used, and 5 repetitions were collected.

### Neural recordings

For the SCT and SRTT extracellular recordings were obtained from the MPC of the monkeys using a system with 7 or 16 independently movable microelectrodes (1-3 MΩ, Uwe Thomas Recording, Germany, S3). Only correct trials were analyzed. All isolated neurons were recorded regardless of their activity during the task, and the recording sites changed from session to session. At each site, raw extracellular membrane potentials were sampled at 40 kHz. Single-unit activity was extracted from these records using the Plexon off-line sorter (Plexon, Dallas, TX). In the present paper we analyzed the activity of 1477 (1074 of Monkey 1 and 403 of Monkey 2) MPC neurons in both monkeys that did not show significant changes in their spontaneous activity during the hold period across all the task (ANOVA, p > 0.05). The functional properties of some of these cells (1083 neurons) have been reported previously [9,10,14]. In addition, using a semichronic, high-density electrode system [23], 26 and 41 MPC cells were recorded simultaneously while monkey M01 was performing the ST and SRTT tasks.

### Neural activation periods

We used the Poisson-train analysis to identify the cell activation periods within each interval defined by two subsequent taps. This analysis determines how improbable it is that the number of action potentials within a specific condition (i.e. target interval and ordinal sequence) was a chance occurrence. For this purpose, the actual number of spikes within a time window was compared with the number of spikes predicted by the Poisson distribution derived from the mean discharge rate during the entire recording of the cell. The measure of improbability was the surprise index (SI) defined as:

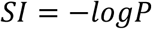

where P was defined by the Poisson equation:

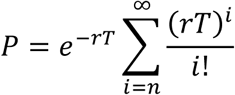

where *P* is the probability that, given the average discharge rate *r*, the spike train for a produced interval *T* contains *n* or more spikes in a trial. Thus, a large *SI* indicates a low probability that a specific elevation in activity was a chance occurrence. This analysis assumes that an activation period is statistically different from the average discharge rate *r*, considering that the firing of the cell is following a non-homogenous Poisson process (see also [49]). The detection of activation periods above randomness has been described previously [4,50]. Importantly, the Poisson-train analysis provided the response-onset latency and the activation period for each cell and for each combination of target interval/serial order.

### Neural trajectories

#### Event time normalization and binarization

We developed a time normalization algorithm to align the neural data from different tapping times of different recording sessions in the same relative time framework. For each neuron, we calculated the produced interval (time between two taps). Then, we subtracted the time of the second tap of a produced interval in the task sequence from all spike and stimulus times (event_times_) and divided them by the produced interval. The tapping times acquired values of minus one and zero, and all the other eventtimes were normalized between these two values. Finally, we added the tap sequence number. Thus, all the normalized values for movement, sensory and spike events acquired values between zero and seven in an SCT trial, as follows:

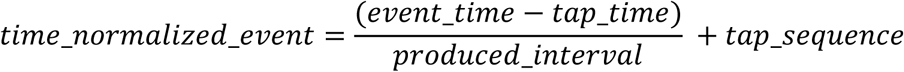

Therefore, the time range of events between the first and the last tap of the normalized data of a trial (UTND) was the same regardless of the target interval. In addition to the trial relative time framework, we also used the target interval normalized data (TIND), which corresponds to the UTND multiplied by the target interval. This time normalization procedure was not necessary for simultaneously recorded data.

#### Trial binarization

For UTND, TIND and simultaneously recorded data, we binarized the neural data by calculating the discharge rate on consecutive windows of 0.02 units. For UTND we always got 50 bins between each pair of taps across target intervals, whereas for TIND and the simultaneously recorded data this number depended on the target interval of the trial. For example, the total number of bins was 23 and 50 for trials with the 450 and 1000 ms intervals, respectively. The binarized data of each neuron was divided by the maximum discharge rate of that particular neuron across all repetitions and target intervals of the SCT.

#### Principal component coefficients matrix

Given a linear transformation of a matrix X into a matrix Y, such that each dimension of Y explains variance of the original data X in descending order, PCA can be described as the search for matrix P that transforms X into Y as follows:

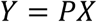

Hence, we first calculated the matrix P using a matrix X that includes all trials and target interval combinations for the visual SCT of our UTND cell population. Using this P on other data guarantees that the same transformation is applied to different neural activity sets. Therefore, using the UTND framework we avoided over- or under-representation of the information for different target intervals, due to the constant total number of bins across conditions.

### Generating neural trajectories

The TIND information for every trial of all neurons constituted the columns of the X’ matrix. The principal component coefficients matrix P were multiplied by the X’ matrix to transform the neural data into the space of the original Y. Using the same transformation matrix for each trial, allowed the comparison of trajectories for different trials and tasks. A locally weighted scatter plot smoothing function was applied to the columns of the Y matrix. The first three dimensions of Y were used to generate graphical three-dimensional trajectories.

### Trajectory radius and variability

The first three PCs explained the 10.7, 3.8 and 2.3 percent of the total variance. The PC1 showed a steep change at the beginning and end of the trial, suggesting a chunking mechanism of the tap sequence in the overall of the neural population-state. In contrast, the PC2 and PC3 showed a strong oscillatory structure with a phase difference of π/2 radians during SCT. For these two PCs, we calculated the centroids of the segments of trajectories between adjacent taps. We measured the radius of the 2D trajectory segment as the mean of the Euclidean distances between the centroid and each point in the trajectory segment. The variability of the trajectory was calculated as the standard deviation of the Euclidean distances between the centroid and each point in the trajectory segment across the six serial order elements (3 of the SC and 3 of the CC) for each target interval. Accordingly, the temporal variability of the behavior for each target interval was computed as the standard deviation of the produced intervals within a trial, namely, the across six serial order elements of the SCT.

### Neural trajectory decoder

We trained a time-delay neural network (TDNN) to decode the produced intervals from the first PC of the simultaneously recorded neural activity during ST. The TDNN architecture had an input layer with 20 time-delays and one hidden 10-unit layer. The output layer consisted of a single unit that was trained to generate a value of 1 when a tap occurred or 0 otherwise. We trained the network using a Bayesian regularization backpropagation algorithm that minimized the mean squared error of the output. The tap time was defined as the time of the peak of the neural network output higher than a threshold of 0.12. We considered a correctly decoded interval when the decoded and the produced taps times difference was less than 60ms. We used 5-fold cross validation to evaluate the performance of the neural network.

### Demixed principal component analysis

The demixed principal component analysis [19] decomposes neural population activity into components capturing the majority of the variance of the data dependent on task parameters. We used the TIND resampled to 30 bins for all target intervals as the input data to the dPCA and the target interval as the marginalization parameter. Therefore, the length of all the trials for all target intervals was the same. We calculated the bin-by-bin Euclidean distance between the 450ms first PC and all the target intervals using the PCA and dPCA analysis.

### Support Vector Machine classifier

We were interested in studying the relation between the neural trajectory dynamics and the instructed interval of the SCT(450ms, 550ms,…, 1000ms). Therefore, we first normalized the length of each segment of the first 8 PCs of the neural trajectory associated to a produced interval (the time between two taps) to 30 bins (see inset Fig. 7A). This step was necessary to avoid a bias associated to the length of the segment. Then, we applied a second layer PCA’ to each of the original neural trajectory segments for each PC independently. We kept the first 3PCs’ as they explained 96% of the variance. As a result, a point in a new three-dimensional coordinate for each 30-bin trajectory segment was obtained (see Fig. 7B). In order to assess which PC had more information about each of the SCT parameters, we carried out a classification procedure for each PC using a Support Vector Machines (SVM) algorithm [51]. Each classifier was retrained 10 times, and we used 5-fold cross validation to evaluate the performance of the classifier. Thus, we identified the PC with more information for each SCT parameter and called it best-PC.

Additionally, we were interested in studying how the size of the neural population used to generate the PCA affected the information contained in the trajectory. We sorted each neuron according to the magnitude of the PCA weights for the best-PC. We iteratively removed the activity of 10% of the neurons with the largest PCA weights for the best-PC until reaching 1% (15 total neurons). Finally, for each population size, we computed the second layer PCAs on the new trajectories and the corresponding SVM classification.

### Oscillatory activity analysis

To characterize the phase, frequency, and amplitude of the neural trajectories, we calculated a series of nonlinear regression models over the residuals of linear regressions on the projected data for the first PC. The general function of the nonlinear models was:

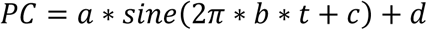

Where *t* is time. In addition, the parameter *a* is the amplitude of the oscillatory function, *b* the frequency, *c* the phase offset, and *c* is a constant. For each trial of both tasks (ST and SRTT) we calculated the mean square error.

### Movement kinematics

We applied the Lucas-Kanade optic flow method to measure the monkey’s arm speed during the ST. This method calculates a flow field from the intensity changes between two consecutive video frames. The analyzed video was recorder with a Microsoft Kinect for Windows camera with a 640×480 resolution. The optic flow method was applied to a smaller area of 140×140 pixels from the original video that contained the monkey’s arm during the whole trial and no other moving objects. The arm’s movement velocity vector was calculated across all frames as the magnitude of the sum of all the individual flow fields vectors which magnitude was larger than a predefined threshold. The velocity vector was calculated from the first to the last tap on each correct trial. We reported the speed as the magnitude of the velocity vector. Posteriorly, the kinematic state of the arm was tagged as movement when the velocity vector was larger than a threshold or dwell otherwise. The tagging algorithm considered a change on the kinematic state when the new state lasted longer than 3 consecutive frames.

### Moving bumps simulations

In order to investigate how the properties of the pattern of neuronal activation affected the generation of population neuronal trajectories. We generated 5 repetitions of simulations of neuronal activity for each target interval. The individual neuronal activation period was composed of the sum of 20 random gamma functions. The activation period was constant for all the neurons on one simulation, but varied with the target interval: 197, 205, 213, 233, and 257ms activation duration for 450, 550, 650, 850, 1000ms target interval respectively. The initial activation time for each neuron was adjusted so that the population activation rate followed a gaussian function as to produce a moving bump pattern. The number of neurons in the simulation was incremented according to the target interval (450ms, 108 neurons; 550ms, 120 neurons; 650ms, 130 neurons; 850ms, 170 neurons; 1000ms, 182 neurons). Fig. 11A shows neurons were added randomly in the intermediate portion of the moving bumps.

## Acknowledgements

We thank Victor de LaFuente, Ranulfo Romo and Roman Rossi for their fruitful comments on the manuscript. We also thank Raul Paulín for his technical assistance. Supported by CONACYT: 236836, CONACYT: 196, and PAPIIT: IN202317 grants to H. Merchant. Jorge Gámez is a doctoral student from Programa de Doctorado en Ciencias Biomédicas, Universidad Nacional Autónoma de México (UNAM) and received fellowship 339118 from CONACYT.

---

**S1 Fig.**
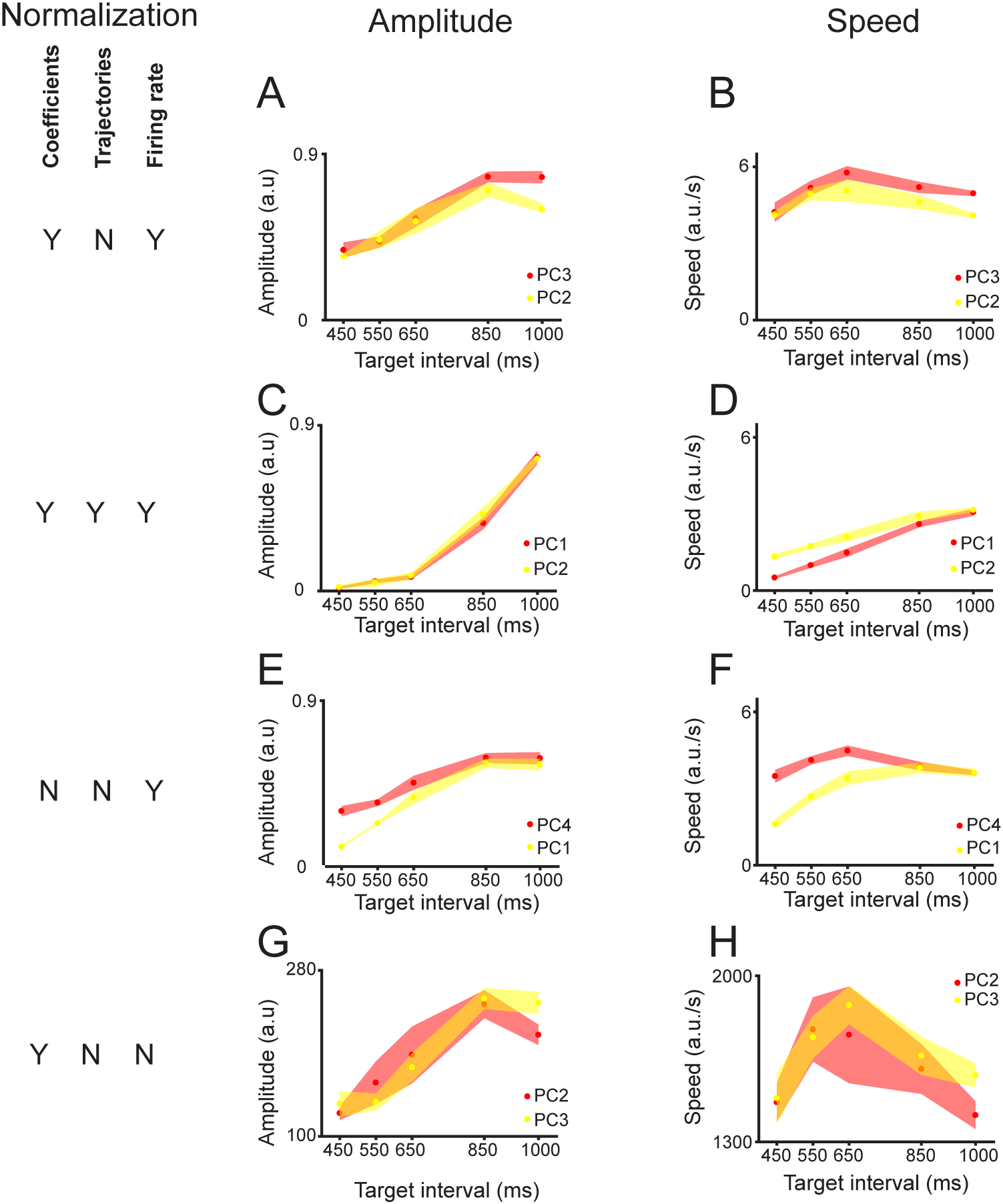
Effect of timing and firing rate normalization on the amplitude and speed of neural trajectories. We used different combinations of the time and firing rate normalization of the neural data in order to calculate the PCA coefficients and then the neural trajectories. We fitted a sine function on each of the first ten PCs and measured their amplitude and speed. For all the possible normalization combinations, we found at least 2 of the first four PCs that showed a robust fit of the sine function that was accompanied by a monotonic increase in the mean and the variability of the trajectory radius and a similar speed across target intervals. Here we show only two PCs (color coded) for each normalization combination (see **A,C,E,G,I**). **(A-F)** were generated using normalized firing rate data to calculate the trajectories. The left row corresponds to PC radial amplitude and the right row to the PC linear speed. **A,B,** coefficients computed with time-normalized but trajectories calculated on actual time bins, as presented across this paper for SCT. (**A**) PC amplitude increased with target interval: PC3, red, data slope= 0.00081, constant = 0.011, R^2^=0.899, p< 0.0001, ANOVA main effect target interval, F(4,20)=128.69,p<0.0001; PC2, yellow, data slope= 0.00053, constant = 0.148, R^2^=0.676, p< 0.0001, ANOVA main effect target interval, F(4,20)=51.54,p<0.0001. (**B**) PC linear speed is similar across target interval: PC3, red, non-significant linear regression, R^2^=0.07, p=0.201, ANOVA main effect target interval, F(4,20)=22.12,p<0.0001; PC2, yellow, non-significant linear regression, R^2^=0.05, p=0.28, ANOVA main effect target interval, F(4,20)=14.36, p<0.0001. **C,D,** coefficients and trajectories are computed using time normalized data. (**C**) PC1, red, data slope= 0.0012, constant = −0.651, R^2^=0.902, p< 0.0001, ANOVA main effect target interval, F(4,20)=875.21,p<0.0001; PC2, yellow, data slope= 0.0013, constant = −0.658, R^2^=0.923, p< 0.0001, ANOVA main effect target interval, F(4,20)=858.7,p<0.0001. (**D**) PC1, red, data slope= 0.0048, constant = −1.638, R^2^=0.98, p<0.0001, ANOVA main effect target interval, F(4,20)=390.94,p<0.0001; PC2, yellow, data slope=0.0034, constant = −0.158, R^2^=0.953, p<0.0001, ANOVA main effect target interval, F(4,20)=160.57, p<0.0001. **E,F,** coefficients and trajectories are computed using actual time data. (**E**) PC4, red, data slope= 0.0005, constant = 0.053, R^2^=0.882, p< 0.0001, ANOVA main effect target interval, F(4,20)=101.86,p<0.0001; PC1, yellow, data slope= 0.00084, constant = −0.225, R^2^=0.899, p< 0.0001, ANOVA main effect target interval, F(4,20)=332.76, p<0.0001. (**F**) PC4, red, non-significant linear regression, R^2^=0.013, p=0.586, ANOVA main effect target interval, F(4,20)=23.35, p<0.0001; PC1, yellow, data slope=0.0034, constant = 0.641, R^2^=0.686, p<0.0001, ANOVA main effect target interval, F(4,20)=100.04, p<0.0001. **G,H**, same as **(A,B)** but using non-normalized firing rate data to calculate the trajectories. (**G**) PC2, red, data slope= 0.175, constant = 62.162, R^2^=0.625, p< 0.0001, ANOVA main effect target interval, F(4,20)=27.58,p<0.0001; PC3, yellow, data slope=0.238, constant = 21.433, R^2^=0.865, p< 0.0001, ANOVA main effect target interval, F(4,20)=101.03, p<0.0001. (**H**) PC2, red, non-significant linear regression, R^2^=0.089, p=0.145, ANOVA main effect target interval, F(4,20)=8.18, p<0.001; PC3, yellow, non-significant linear regression, R^2^=0.0002, p=0.939, ANOVA main effect target interval, F(4,20)=16.37, p<0.0001.

**S2 Fig.**
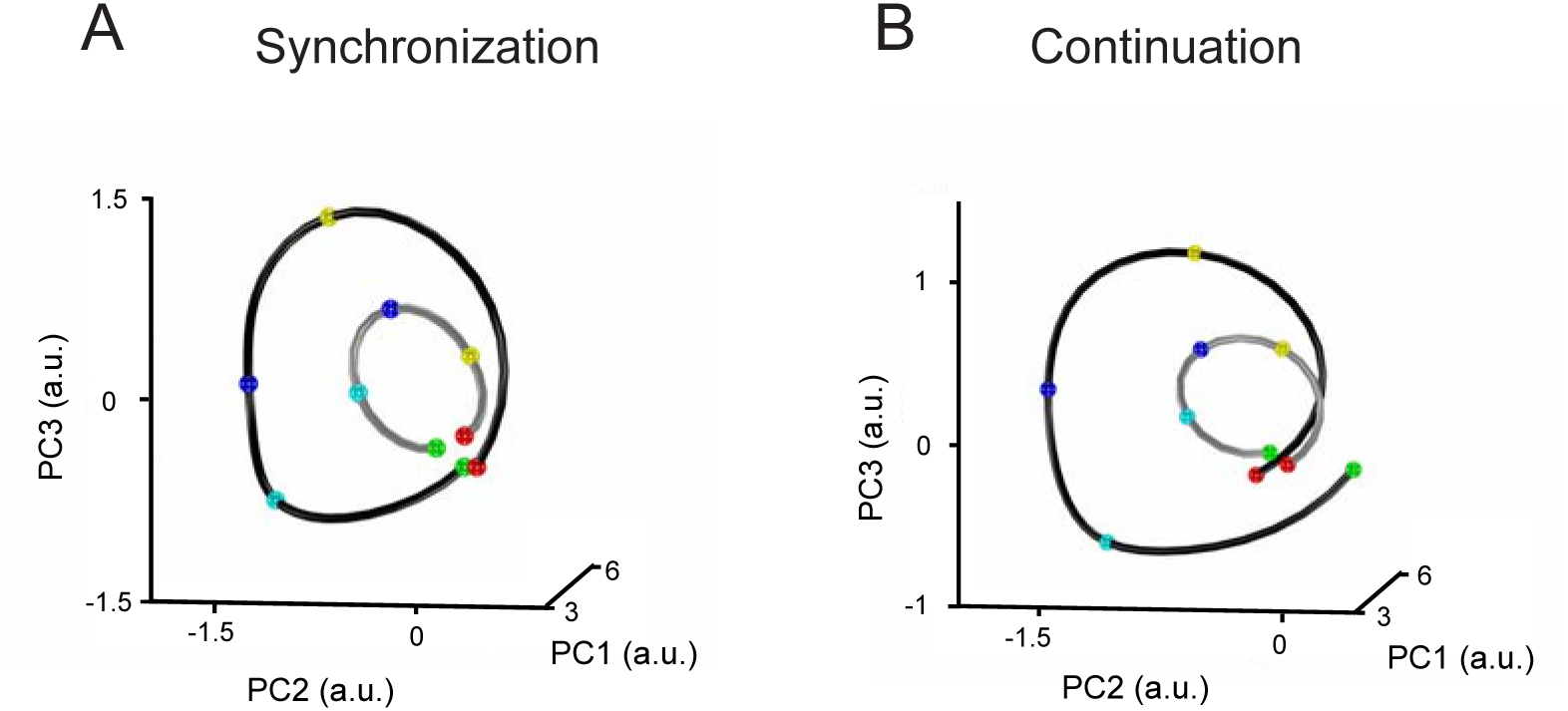
State trajectory progress during SCT. **A,B,** One trajectory loop for the second produced interval of the (A) SC and (B) CC, during a 450ms (light gray) and a 1000ms (dark gray) target interval. Trajectory progression marked as colored spheres: previous tap (green), 1^st^ inter-tap quarter (cyan), 2^nd^ inter-tap quarter/half interval (blue), 3^rd^ inter-tap quarter (yellow), and next tap (red). Therefore, the neural trajectories follow circular paths with different radii that increase according to the target interval but with similar speed profiles.

**S3 Fig.**
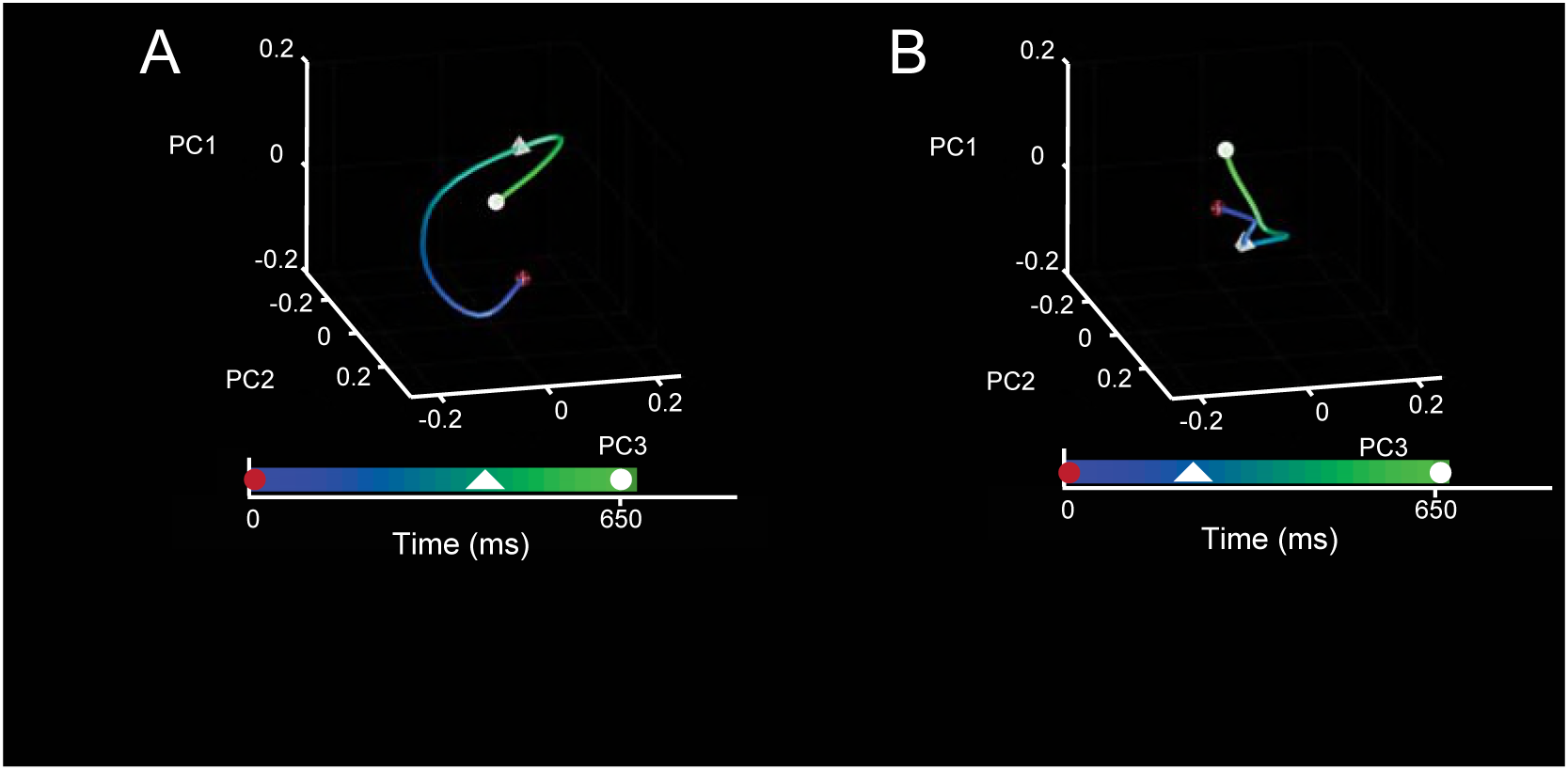
State trajectories during ST and SRTT using simultaneously recorded neurons. **A,B,** Three dimensional neural dynamics trajectory of a 650ms single ST (A) and SRTT (B) intervals. Elapsed time is colorcoded. The previous and the next taps are marked as red and white spheres respectively. The stimuli are marked as a white pyramid.

## References

1. Patel AD. The Evolutionary Biology of Musical Rhythm: Was Darwin Wrong? PLoS Biol. 2014;12: 1–6. doi:10.1371/journal.pbio.1001821

2. Teki S, Grube M, Kumar S, Griffiths TD. Distinct neural substrates of duration-based and beat-based auditory timing. J Neurosci. 2011;31: 3805–3812. doi:10.1523/JNEUROSCI.5561-10.2011

3. Grahn JA. Neuroscientific Investigations of Musical Rhythm: Recent Advances and Future Challenges. Contemp Music Rev. 2009;28: 251–277. doi:10.1080/07494460903404360

4. Merchant H, Pérez O, Bartolo R, Méndez JC, Mendoza G, Gámez J, et al. Sensorimotor neural dynamics during isochronous tapping in the medial premotor cortex of the macaque. Eur J Neurosci. 2015;41: 586–602. doi:10.1111/ejn.12811

5. Merchant H, Georgopoulos AP. Neurophysiology of Perceptual and Motor Aspects of Interception. J Neurophysiol. 2006;95: 1–13. doi:10.1152/jn.00422.2005

6. Kotz SAE, Schwartze M. Differential Input of the Supplementary Motor Area to a Dedicated Temporal Processing Network: Functional and Clinical Implications. Front Integr Neurosci. 2011;5. doi:10.3389/fnint.2011.00086

7. Merchant H, Grahn J, Trainor L, Rohrmeier M, Fitch WT. Finding the beat: a neural perspective across humans and non-human primates. Philos Trans R Soc Lond B Biol Sci. 2015;370: 20140093. doi:10.1098/rstb.2014.0093

8. Merchant H, Harrington DL, Meck WH. Neural basis of the perception and estimation of time. Annu Rev Neurosci. 2013;36: 313–336. doi:10.1146/annurev-neuro-062012-170349

9. Merchant H, Pérez O, Zarco W, Gámez J. Interval tuning in the primate medial premotor cortex as a general timing mechanism. J Neurosci. 2013;33: 9082–9096. doi:10.1523/JNEUROSCI.5513-12.2013

10. Crowe DA, Zarco W, Bartolo R, Merchant H. Dynamic Representation of the Temporal and Sequential Structure of Rhythmic Movements in the Primate Medial Premotor Cortex. J Neurosci. 2014;34: 11972–11983. doi:10.1523/JNEUROSCI.2177-14.2014

11. Bartolo R, Prado L, Merchant H. Information Processing in the Primate Basal Ganglia during Sensory-Guided and Internally Driven Rhythmic Tapping. J Neurosci. 2014;34: 3910–3923. doi:10.1523/JNEUROSCI.2679-13.2014

12. Mello GBM, Soares S, Paton JJ. A Scalable Population Code for Time in the Striatum. Curr Biol. Elsevier Ltd; 2015;25: 1113–1122. doi:10.1016/j.cub.2015.02.036

13. Wang J, Narain D, Hosseini EA, Jazayeri M. Flexible timing by temporal scaling of cortical responses. Nat Neurosci. Springer US; 2018;21: 102–110. doi:10.1038/s41593-017-0028-6

14. Merchant H, Zarco W, Pérez O, Prado L, Bartolo R. Measuring time with different neural chronometers during a synchronization-continuation task. Proc Natl Acad Sci U S A. 2011;108: 19784–19789. doi:10.1073/pnas.1112933108

15. Knudsen EB, Powers ME, Moxon KA. Dissociating Movement from Movement Timing in the Rat Primary Motor Cortex. J Neurosci. 2014;34: 15576–15586. doi:10.1523/JNEUROSCI.1816-14.2014

16. Jazayeri M, Shadlen MN. A Neural Mechanism for Sensing and Reproducing a Time Interval. Curr Biol. 2015;25: 2599–2609. doi:10.1016/j.cub.2015.08.038

17. Merchant H, Bartolo R. Primate beta oscillations and rhythmic behaviors. J Neural Transm. 2018;125: 461–470. doi:10.1007/s00702-017-1716-9

18. Cunningham JP, Yu BM. Dimensionality reduction for large-scale neural recordings. Nat Neurosci. 2014;17: 1500–1509. doi:10.1038/nn.3776

19. Kobak D, Brendel W, Constantinidis C, Feierstein CE, Kepecs A, Mainen ZF, et al. Demixed principal component analysis of neural population data. Elife. 2016;5: 1–36. doi:10.7554/eLife.10989

20. Murray JM, Escola GS. Learning multiple variable-speed sequences in striatum via cortical tutoring. Elife. 2017;6: 1–24. doi:10.7554/eLife.26084

21. Rossi-Pool R, Zainos A, Alvarez M, Zizumbo J, Vergara J, Romo R. Decoding a Decision Process in the Neuronal Population of Dorsal Premotor Cortex. Neuron. 2017;96: 1432–1446.e7. doi:10.1016/j.neuron.2017.11.023

22. Remington ED, Narain D, Hosseini EA, Jazayeri M. Flexible Sensorimotor Computations through Rapid Reconfiguration of Cortical Dynamics. Neuron. Elsevier Inc.; 2018;98: 1005–1019.e5. doi:10.1016/j.neuron.2018.05.020

23. Mendoza G, Peyrache A, Gámez J, Prado L, Buzsáki G, Merchant H. Recording extracellular neural activity in the behaving monkey using a semi-chronic and high-density electrode system. J Neurophysiol. 2016;116: 563–574. doi:10.1152/jn.00116.2016

24. Donnet S, Bartolo R, Fernandes JM, Cunha JPS, Prado L, Merchant H. Monkeys time their pauses of movement and not their movement-kinematics during a synchronization-continuation rhythmic task. Journal of Neurophysiology. 2014. doi:10.1152/jn.00802.2013

25. Gámez J, Yc K, Ayala YA, Dotov D, Prado L, Merchant H. Predictive rhythmic tapping to isochronous and tempo changing metronomes in the nonhuman primate. Ann N Y Acad Sci. 2018; 1–20. doi:10.1111/nyas.13671

26. Hardy NF, Buonomano D V. Encoding Time in Feedforward Trajectories of a Recurrent Neural Network Model. Neural Comput. 2018;30: 378–396. doi:10.1162/neco_a_01041

27. Goudar V, Buonomano D V. Encoding sensory and motor patterns as time-invariant trajectories in recurrent neural networks. Elife. 2018;7: 1–28. doi:10.7554/eLife.31134

28. Mendoza G, Merchant H. Motor system evolution and the emergence of high cognitive functions. Prog Neurobiol. 2014;122: 73–93. doi:10.1016/j.pneurobio.2014.09.001

29. Churchland MM, Cunningham JP, Kaufman MT, Foster JD, Nuyujukian P, Ryu SI, et al. Neural population dynamics during reaching. Nature. Nature Publishing Group; 2012;487: 51–56. doi:10.1038/nature11129

30. Russo AA, Bittner SR, Perkins SM, Seely JS, London BM, Lara AH, et al. Motor Cortex Embeds Muscle-like Commands in an Untangled Population Response. Neuron. Elsevier Inc.; 2018;97: 953–966.e8. doi:10.1016/j.neuron.2018.01.004

31. Karmarkar UR, Buonomano D V. Timing in the absence of clocks: encoding time in neural network states. Neuron. 2007;53: 427–438. doi:10.1016/j.neuron.2007.01.006

32. Merchant H, Yarrow K. How the motor system both encodes and influences our sense of time. Curr Opin Behav Sci. Elsevier Ltd; 2016;8: 22–27. doi:10.1016/j.cobeha.2016.01.006

33. Paton JJ, Buonomano D V. The Neural Basis of Timing: Distributed Mechanisms for Diverse Functions. Neuron. Elsevier Inc.; 2018;98: 687–705. doi:10.1016/j.neuron.2018.03.045

34. Merchant H, Bartolo R, Pérez O, Méndez JC, Mendoza G, Gámez J, et al. Neurophysiology of Timing in the Hundreds of Milliseconds: Multiple Layers of Neuronal Clocks in the Medial Premotor Areas. In: Merchant H, de Lafuente V, editors. Neurobiology of Interval Timing. New York, NY: Springer New York; 2014. pp. 143–154. doi:10.1007/978-1-4939-1782-2_8

35. Gibbon J, Malapani C, Dale CL, Gallistel CR. Toward a neurobiology of temporal cognition: Advances and challenges. Curr Opin Neurobiol. 1997;7: 170–184. doi:10.1016/S0959-4388(97)80005-0

36. Merchant H, Zarco W, Prado L. Do we have a common mechanism for measuring time in the hundreds of millisecond range? Evidence from multiple-interval timing tasks. J Neurophysiol. 2008;99: 939–949. doi:10.1152/jn.01225.2007

37. García-Garibay O, Cadena-Valencia J, Merchant H, de Lafuente V. Monkeys Share the Human Ability to Internally Maintain a Temporal Rhythm. Front Psychol. 2016;7: 1–12. doi:10.3389/fpsyg.2016.01971

38. Mendez JC, Prado L, Mendoza G, Merchant H. Temporal and Spatial Categorization in Human and Non-Human Primates. Front Integr Neurosci. 2011;5: 1–10. doi:10.3389/fnint.2011.00050

39. Simen P, Balci F, deSouza L, Cohen JD, Holmes P. A Model of Interval Timing by Neural Integration. J Neurosci. 2011;31: 9238–9253. doi:10.1523/JNEUROSCI.3121-10.2011

40. Merchant H, Averbeck BB. The Computational and Neural Basis of Rhythmic Timing in Medial Premotor Cortex. J Neurosci. 2017;37: 4552–4564. doi:10.1523/JNEUROSCI.0367-17.2017

41. Pérez O, Merchant H. The synaptic properties of cells define the hallmarks of interval timing in a recurrent neural network. The Journal of Neuroscience. 2018. doi:10.1523/JNEUROSCI.2651-17.2018

42. Kaufman MT, Churchland MM, Ryu SI, Shenoy K V. Cortical activity in the null space: permitting preparation without movement. Nat Neurosci. Nature Publishing Group; 2014;17: 440–448. doi:10.1038/nn.3643

43. Crowe DA, Averbeck BB, Chafee M V. Rapid Sequences of Population Activity Patterns Dynamically Encode Task-Critical Spatial Information in Parietal Cortex. J Neurosci. 2010;30: 11640–11653. doi:10.1523/JNEUROSCI.0954-10.2010

44. Jin DZ, Fujii N, Graybiel AM. Neural representation of time in cortico-basal ganglia circuits. Proc Natl Acad Sci. 2009;106: 19156–19161. doi:10.1073/pnas.0909881106

45. Gouvêa TS, Monteiro T, Motiwala A, Soares S, Machens C, Paton JJ. Striatal dynamics explain duration judgments. Elife. 2015;4: 1–14. doi:10.7554/eLife.11386

46. Pastalkova E, Itskov V, Amarasingham A, Buzsaki G. Internally Generated Cell Assembly Sequences in the Rat Hippocampus. Science (80-). 2008;321: 1322–1327. doi:10.1126/science.1159775

47. MacDonald CJ, Lepage KQ, Eden UT, Eichenbaum H. Hippocampal “time cells” bridge the gap in memory for discontiguous events. Neuron. Elsevier Inc.; 2011;71: 737–749. doi:10.1016/j.neuron.2011.07.012

48. Zarco W, Merchant H, Prado L, Mendez JC. Subsecond timing in primates: comparison of interval production between human subjects and rhesus monkeys. J Neurophysiol. 2009;102: 3191–202. doi:10.1152/jn.00066.2009

49. Perez O, Kass RE, Merchant H. Trial time warping to discriminate stimulus-related from movement-related neural activity. J Neurosci Methods. Elsevier B.V.; 2013;212: 203–210. doi:10.1016/j.jneumeth.2012.10.019

50. Merchant H, Battaglia-mayer A. Effects of optic flow in motor cortex and area 7a Effects of Optic Flow in Motor Cortex and Area 7a. 2001;

51. Cortes C, Vapnik V. Support-vector networks. Mach Learn. 1995;20: 273–297. doi:10.1007/BF00994018

